# The E2 SUMO-conjugating enzyme UBE2I coordinates the oocyte and zygotic transcriptional programs

**DOI:** 10.1101/2022.12.06.519314

**Authors:** Shawn M. Briley, Avery A. Ahmed, Peixin Jiang, Sean M. Hartig, Karen Schindler, Stephanie A. Pangas

**Author notes:** Corresponding Author: Stephanie A. Pangas, PhD, One Baylor Plaza, Baylor College of Medicine, Houston, TX 77030.

## Abstract

In mammals, meiotically competent oocytes develop cyclically during ovarian folliculogenesis. During folliculogenesis, prophase I arrested oocytes are transcriptionally active, producing and storing transcripts required for their growth and for early stages of embryogenesis prior to the maternal to zygotic transition. Defective oocyte development during folliculogenesis leads to meiotic defects, aneuploidy, follicular atresia, or non-viable embryos. Here we generated a novel oocyte-specific knockout of the SUMO E2 ligase, *Ube2i*, using *Zp3-cre* to test its function during folliculogenesis.

*Ube2i Zp3-cre*+ female mice are sterile with oocytes that arrest in meiosis I with defective spindles and chromosome alignment. Fully grown mutant oocytes abnormally maintain transcription but downregulate maternal effect genes and prematurely activate the zygotic transcriptional program. Thus, this work uncovers UBE2i as a novel orchestrator of chromatin and transcriptional regulation in mouse oocytes.

**Teaser:** Oocyte-specific deletion of *Ube2i* causes loss of transcriptional repression and premature activation of the zygotic genome.

## Introduction

In mammals, oocyte development begins embryonically but is not completed until the oocyte is fertilized following ovulation in adults. Prior to birth, mouse oocytes begin meiosis at embryonic day (E13.5) and proceed through the early stages of prophase I to arrest in the diplotene stage (*1, 2*). Oocytes remain arrested in prophase I while stored as primordial follicles and following recruitment into the growing pool of follicles, where they develop to fully-grown oocytes capable of resuming meiosis, *i.e.*, meiotic competence. During folliculogenesis, prophase I oocytes are transcriptionally active, and produce and store transcripts needed for meiotic maturation and early embryogenesis; once fully grown, transcription is silenced and the chromatin undergoes structural changes within the oocyte nucleus, also called the germinal vesicle (GV) (*3, 4*). Following the midcycle surge of luteinizing hormone (LH), oocytes resume meiosis, the nuclear envelope breaks down (GVBD), and the process of spindle assembly begins. Oocytes proceed through pro-metaphase to reach metaphase I and must satisfy the spindle assembly checkpoint (SAC) before homologous chromosomes are separated and the first polar body is extruded, marking the end of meiosis I. A second meiotic arrest then occurs at metaphase II, where the oocytes, from this point typically referred to as eggs, remain arrested until fertilization (*5*). Oocytes remain transcriptionally silent throughout meiotic maturation, and transcription does not resume until after fertilization when zygotic genome activation (ZGA) occurs during the maternal-to-zygotic transition (MZT) (*6*).

SUMOylation is an essential covalent post-translational protein modification (PTM) whereby a small ubiquitin-like modifier protein (SUMO) is attached to a lysine of the target protein with a specific sequence (ψ-K-x-E where ψ is a hydrophobic amino acid, K is lysine, x is arbitrary, and E is glutamic acid). This modification is reversible and highly dynamic, making detection of endogenously modified proteins challenging (*7*). There are four SUMO protein isoforms in mammals (SUMO1-4), three of which, SUMO1, SUMO2, and SUMO3, are expressed in mouse oocytes (*8, 9*). SUMOylation uses a stepwise enzymatic cascade (E1 “activating”, E2 “conjugating’, and E3 “ligase” enzymes) similar to ubiquitination. However, unlike ubiquitination, in which there are numerous genes encoding the E2 conjugating enzymes, there is only one known E2 in the SUMOylation pathway, UBE2I (also known as UBC9), and only a handful of E3 ligases, which are not required for SUMOylation (*7, 10*). Mice lacking *Ube2i* die at the early post-implantation stage, with defects in chromosome condensation and segregation with a range of nuclear structural defects (*11*).

Protein SUMOylation impacts diverse functions, including protein-protein interaction, subcellular localization, proteostasis, and transcription (*12*). SUMOylation predominantly targets nuclear proteins, and regulates DNA replication, DNA damage repair, and chromatin (*13–17*).

SUMOylation also maintains cell identity and pluripotency (*12*)in mouse embryonic stem cells (ESC (*13*). In addition, SUMO modification of repressive chromatin complexes in ESCs prevents chromatin from opening (*14*). SUMOylation is also an essential regulator of meiosis (*18–20*) with functions identified in synaptonemal complex disassembly, suppression of nondisjunction, spindle formation, and chromosome congression and separation. Taken together, these studies identify SUMOylation as a key regulator of meiosis, chromatin configuration, and transcriptional activity in many model systems, though surprisingly, this function is less studied in mammalian oocyte biology. Increasing evidence indicates SUMOylation has critical but mostly uncharacterized roles in mouse oocyte biology, including regulation of meiotic resumption, meiotic progression, and spindle integrity (*9, 18, 21-23*) We previously generated a mouse strain with conditional deletion of *Ube2i* in oocytes beginning at the primordial follicle stage (*i.e.,* in the resting pool and prior to the start of folliculogenesis) (*18*).

Female mice were sterile with premature depletion of ovarian follicles in early adulthood reminiscent of primary ovarian insufficiency. *Ube2i* deletion in oocytes of primordial follicles additionally caused depletion of growing follicles, suggesting that oocyte SUMOylation is required for the stability of both non-growing and growing follicles (*18*). To test this hypothesis, we generated a novel *Ube2i* oocyte knockout using *Zp3-cre*, an oocyte-specific cre recombinase line that deletes *loxP*-flanked (‘floxed’) alleles in growing follicles (*24*) and thus should not directly affect the non-growing pool of primordial follicles in the ovary. Here, we show that *Ube2i Zp3-cre+* female mice are sterile but have a stable ovarian reserve and growing follicle pool. Unexpectedly, analysis of *Ube2i Zp3-cre+* mice uncovered additional roles for UBE2I in mouse oocytes that included establishing a fully condensed chromatin configuration, regulating global transcriptional silencing, and preventing premature activation of zygotic genome activation genes. Thus, our approach uncovered SUMOylation as a major regulatory process that orchestrates transcriptional silencing at the end of oocyte growth and subsequent timing of zygotic genome activation in mouse oocytes.

## Results

### Oocyte-specific deletion of *Ube2i* at the primary follicle stage causes female sterility but maintains the ovarian reserve

To determine the role of UBE2I in growing oocytes, male mice carrying cre-recombinase under control of the regulatory sequences of the mouse zona pellucida 3 (*Zp3*) gene (*Zp3-cre*) were crossed to *Ube2i^loxP/loxP^*female mice to specifically delete *Ube2i* in oocytes within primary follicles (Fig. 1A) (*24*). Loss of *Ube2i* was confirmed by quantitative reverse transcription PCR (qRT-PCR) of fully grown oocytes from antral follicles, demonstrating *Ube2i* transcript expression in *Ube2i^loxP/loxP^ Zp3-cre+* (*Ube2i Zp3-cre+*) oocytes was reduced by 99.4% compared to the *Ube2i^loxP/loPx^* (control) littermates (P<0.0001) (Fig. 1B). To determine if loss of *Ube2i* affected fertility, six-week-old female control or *Ube2i Zp3-cre+* female littermates were pair housed and continually mated with sexually mature wild type males for six months. While control females produced an average of 6.4 ± 0.5 pups per litter, *Ube2i Zp3-cre+* females failed to produce any liters, demonstrating that oocyte-specific deletion of *Ube2i*using *Zp3-cre* causes female sterility (Fig. 1C, P=0.0003).

**Fig. 1.**
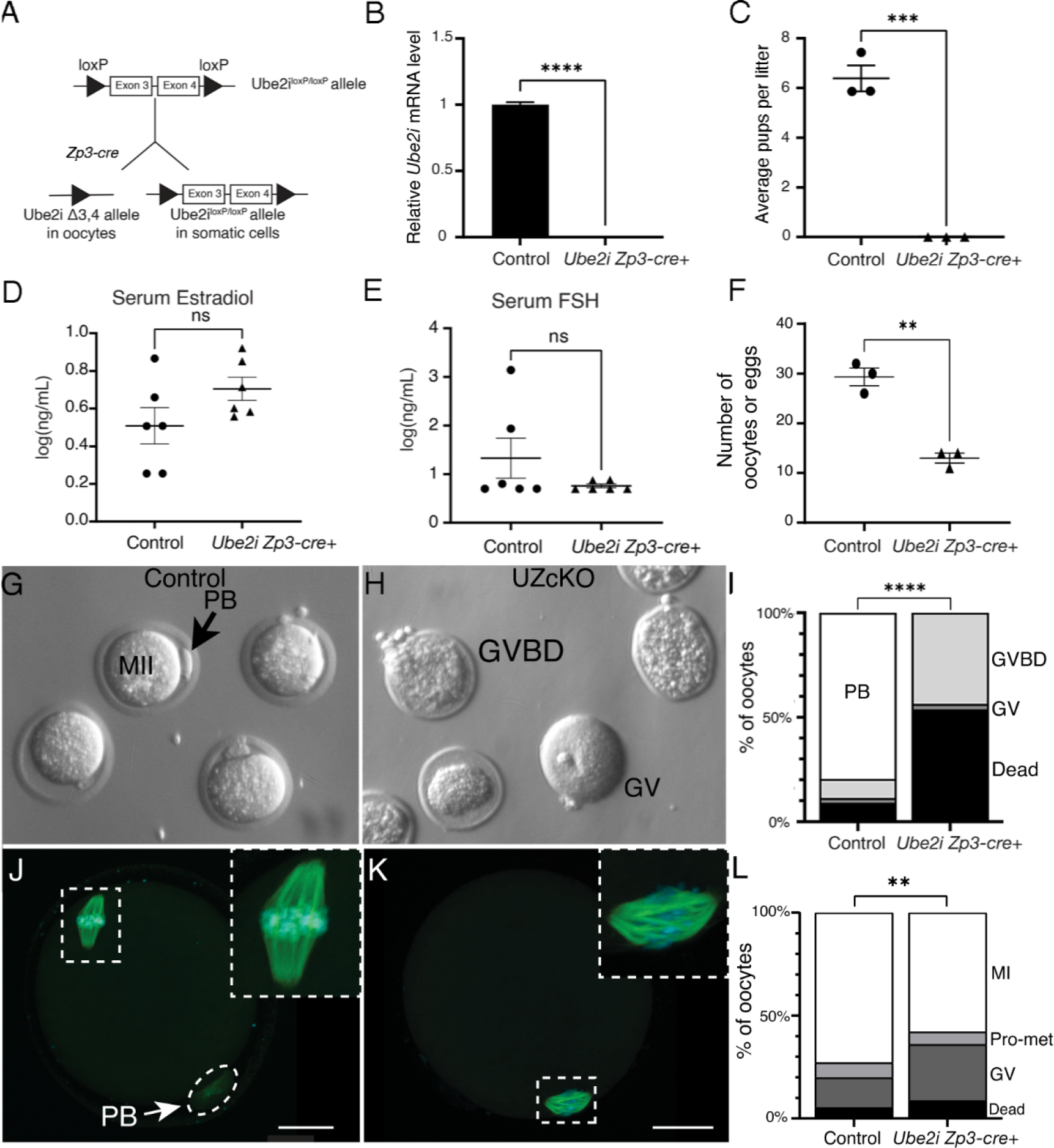
Deletion of *Ube2i^loxP/loxP^* using *Zp3-cre* causes female sterility. A) Schematic for generating the *Ube2i^loxP/loxP^Zp3-Cre+* mice model. B) qRT-PCR analysis for *Ube2i* in *Ube2i Zp3-cre+* GV oocytes (three pools of n=86 from *Ube2i* Zp3-cre+ or 97 control oocytes from 3 animals per genotype), normalized to *Gapd* and relative to control *Ube2i* mRNA levels shows successful loss of *Ube2i* (P < 0.0001 by unpaired t-test) with mean +/- s.e.m shown. C) Six-week-old control or *Ube2i Zp3-cre+* females were continuously housed with fertile wild type males for six months and the presence of newborn pups monitored daily. Control females averaged liter sizes of 6.4 +/- 0.5 pups per liter while *Ube2i Zp3-cre+* females were sterile (P = 0.0003, by unpaired t-test. n = 3 for both genotypes). Serum levels of estradiol (D) and FSH (E) were similar between genotypes (n=6 per genotype) as measured by ELISA (E2, P = 0.1171 unpaired t-test; FSH, P = 0.2269 Welch’s t-test; statistical analysis on log transformed data). F) By superovulation, fewer oocytes are collected from oviduct ampulla in *Ube2i Zp3-cre* females than control littermates (P<0.05 by unpaired t-test), n=3 mice per genotype). (G) Eggs ovulated from control mice contain a polar body, indicating they are arrested at metaphase II. (H) Oocytes ovulated from *Ube2i Zp3-cre+* females lack a polar body. (I) Ovulated oocytes collected from *Ube2i Zp3-cre+* females are significantly more likely to lack a polar body or be dead than ovulated oocytes collected from control females (P <0.0001 by chi-square test. Control n = 88, *Ube2i Zp3-cre+* n = 39). Control (J) and *Ube2i Zp3-cre+* (K) oocytes collected after superovulation and immunostained with-tubulin (green) and Hoechst (DNA) (blue), confocal z-stack projection, scale bars are 20µm. (L) α In vitro culture of GV oocytes isolated from control and *Ube2i Zp3-cre+* shows impaired meiotic resumption and progression over 8.5hr in (P = 0.0077 by chi-square test. Control n = 151, *Ube2i Zp3- cre+* n = 114). In panels C-F, individual animals are show by black circles (control) or black triangles (*Ube2i Zp3-cre+*) with mean +/- s.e.m shown.

Our previous oocyte-specific knockout, *Ube2i Gdf9-icre+,* which begins recombination at the primordial follicle stage (*25*), shows premature depletion of the ovarian reserve with almost complete depletion of all ovarian follicles by 6 months of age (*18*). To assess if loss of *Ube2i* in oocytes of growing follicles also affected stability of the ovarian reserve, ovaries from six-month control and *Ube2i Zp3-cre+* females were collected and ovarian follicle morphometrics determined. Unlike 6-month old *Ube2i^loxP/loxP^Gdf9-icre* ovaries (*18*), *Ube2i Zp3-cre+* ovaries were a similar size and contained similar numbers of all follicle stages compared to control littermates, except for a significant decrease in the number of primary follicles, the stage at which *Zp3-cre* is first expressed (Fig. S1) (*18*). In line with maintenance of the ovarian reserve, *Ube2i Zp3-cre+* female mice had similar levels of estradiol (E2) and follicle stimulating hormone (FSH) as their littermate controls (Fig. 1 D, E). These results indicate that although loss of *Ube2i* in growing oocytes causes sterility, it minimally disrupts follicle dynamics and ovarian hormonal feedback.

### *Ube2i Zp3-cre+* oocytes arrest at metaphase I

The presence of corpora lutea in histologic sections of ovaries of *Ube2i Zp3-cre+* (Fig. S1) indicated these mice potentially ovulate despite being sterile. Therefore, ovulation was assessed using stimulation by exogenous gonadotropin injection (“superovulation”). Compared to controls, *Ube2i Zp3- cre+* females ovulated significantly fewer metaphase II (MII) eggs (P= 0.0138) compared to control females (Fig. 1F). Within the oviductal ampulla, eggs from control females were enclosed in an expanded cumulous-oocyte-complex (COC), whereas those ovulated from *Ube2i Zp3-cre+* females lacked cumulus cells around the oocyte (data not shown). In addition, ovulated cells from control mice were at the correct MII stage with visible polar bodies as expected; however, ovulated oocytes from *Ube2i Zp3-cre+* females lacked a visible polar body (Fig. 1G, H). To verify meiotic stage of the ovulated cells, we analyzed spindle shape and configuration via alpha-tubulin immunostaining and DNA labeling. Control ovulated eggs contained an MII spindle, whereas *Ube2i Zp3-cre+* oocytes arrested in metaphase I (MI), with disorganized spindles and misnaligned chromosomes (Fig. 1J, K).

To examine MI in more detail, we determined the ability of fully grown, germinal vesicle (GV) stage oocytes to undergo meiotic resumption and progression *in vitro*. Fully grown GV oocytes were harvested from control and *Ube2i Zp3-cre+* ovaries following hormone stimulation into medium containing 2.5µM milrinone, which maintains meiotic arrest. After removal of milrinone, oocytes were cultured for 8.5 hours, fixed, and analyzed for GVBD and MI progression. The progression of oocytes from the GV stage to MI was significantly altered between control and *Ube2i ZP3-cre+*oocytes (P=0.0077) (Fig. 1L). Similar percentages of oocytes died (control 5.3*%, Ube2i Zp3-cre*+ 8.8%) or reached pro-metaphase (control 7.3*%, Ube2i Zp3-cre*+ 6.1%). However, the percentage of *Ube2i Zp3- cre*+ oocytes that remained at the GV stage (27.2%) was nearly double that of control oocytes (14.6%), and control oocytes were also more successful at reaching MI (72.8%) than *Ube2i Zp3-cre*+ oocytes (57.9%) (Fig. 1L). These data indicate that *Ube2i Zp3-cre*+ oocytes have reduced meiotic competence compared to control oocytes, supporting the MI arrest phenotype observed in oocytes ovulated from *Ube2i Zp3-cre*+ mice.

MI oocytes were further analyzed for spindle architecture and chromosome alignment by confocal microscopy for microtubules using α-tubulin antibodies and Hoechst to detect DNA (Fig. 2A- F). For classification, oocytes with multiple chromosomes lagging or not aligned on the metaphase plate were classified as having “poor” alignment, while oocytes with all or nearly all chromosomes aligned at the metaphase plate were classified as having “good” alignment. A significant majority (60.6%, P < 0.0001) of *Ube2i Zp3-cre+* MI oocytes had poorly aligned chromosomes, in contrast to “good” chromosome alignment in the majority (81.2%) of control MI oocytes (Fig. 2). In addition to misaligned chromosomes, *Ube2i Zp3-cre+* oocytes often displayed abnormal spindles, including those that appear stretched, shortened, or disorganized (Fig. 2D-F). To quantify differences in spindle length and width, MI spindles were measured from pole to pole and at the widest point of the mid spindle. Compared to controls, *Ube2i Zp3-cre+* spindles were found to be significantly shorter (P=0.0444) and narrower (P=0.0285) than control spindles (Fig. 2H, 2I). Chromosome spreads from MI oocytes showed that *Ube2i Zp3-cre*+ oocytes successfully form bivalents; however, they appeared less condensed compared to control bivalents (Fig. 2J, 2K).

**Fig. 2.**
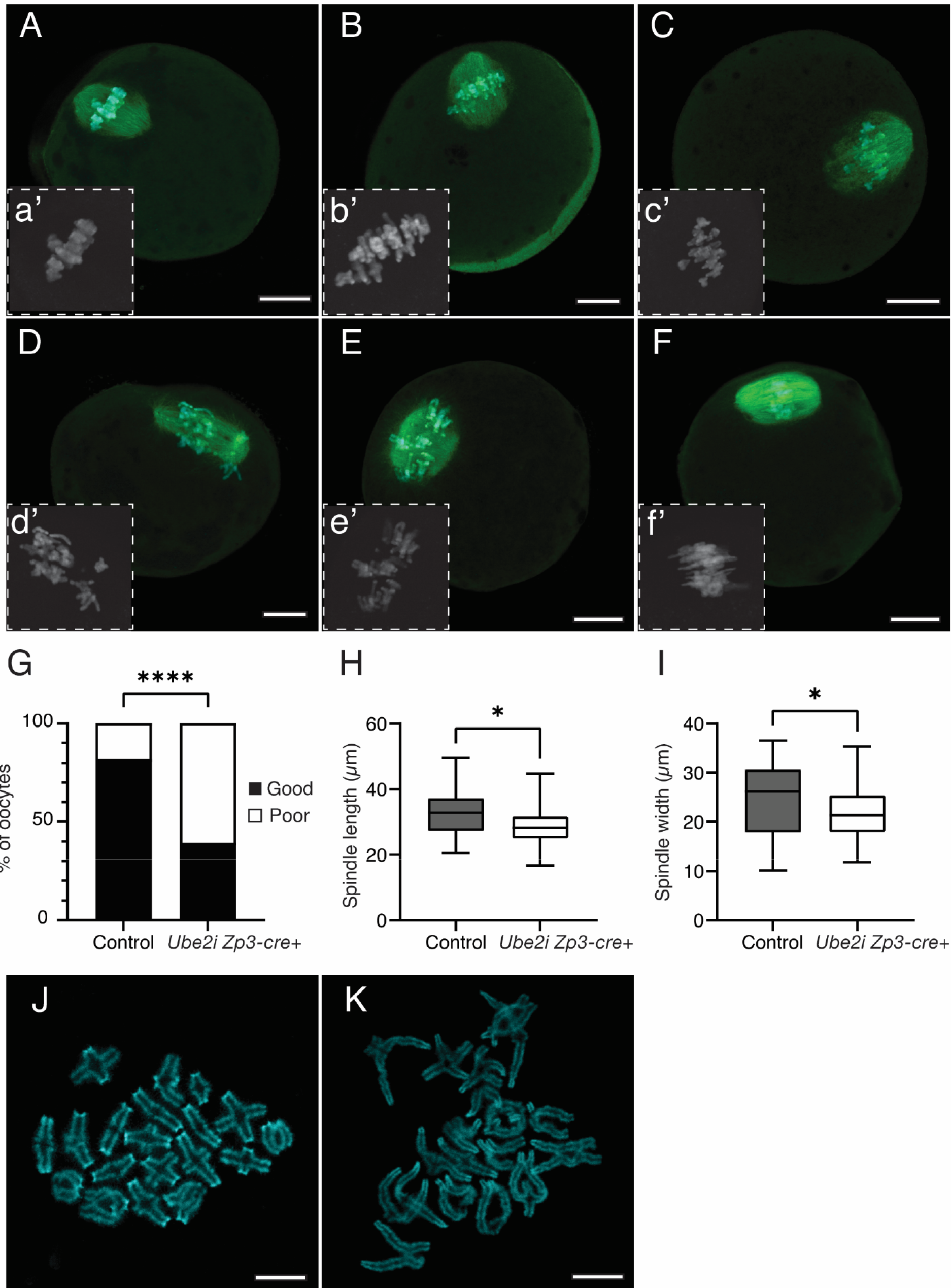
*Ube2i Zp3-cre+* oocytes have misaligned chromosomes at metaphase I. Oocytes were cultured in CZB medium supplemented with glutamine for 8.5 hours then immunostained with anti-alpha-tubulin (green) antibody and Hoechst, and chromosome alignment was analyzed. (A-C) representative images of control oocyte metaphase I spindles, insets (a’-c’) showing chromosomes by Hoechst staining (D-F) Representative images of *Ube2i Zp3-cre+* metaphase I spindles, insets (d’-f’) showing chromosomes by Hoechst staining. Representative confocal z-slices, all scale bars are 20µm. G) Quantification of chromosome alignment as determined by chromosome localization on the metaphase plate. ****P < 0.001, Fisher’s exact test. Control n = 110, *Ube2i Zp3-cre+* n = 66. H) Metaphase I spindle length from pole to pole of the bivalent spindle *P=0.0444, unpaired t-test. Control n = 47, *Ube2i Zp3-cre+* n = 35. I) Metaphase I spindle width measured at the widest point of the bivalent spindle. *P=0.0285, unpaired t-test. Control n = 47, *Ube2i Zp3-cre+*n = 35. J) Chromosome spreads showing MI bivalents from control metaphase I oocytes. K) Chromosome spreads showing less condensed MI bivalents from *Ube2i Zp3-cre+* metaphase I oocytes. Scale bars in J and K are 10µm.

Because of the misaligned chromosomes, we further analyzed if microtubule attachment to kinetochores was disrupted in control and *Ube2i Zp3-cre+* oocytes (Fig. 3). We used a cold-stable assay as low temperatures depolymerize microtubules not attached to kinetochores, while microtubules attached to kinetochores are relatively stable (*26*). GV oocytes were cultured to mid-MI (7 hours) then placed in ice-cold MEM prior to fixation and analyzed for microtubules (α-tubulin) and kinetochores (CREST) staining. Due to the abnormal chromosome structure and poor alignment in *Ube2i Zp3-cre+* oocytes at MI (Fig. 2), it was not possible to determine the precise number of kinetochore-microtubule attachments, so only the presence or absence of cold-stable microtubules was scored. Control oocytes were significantly more likely to have cold-stable microtubules (97.2%) than *Ube2i Zp3-cre+* oocytes (79.2%, P=0.0333) (Fig. 3A-C). Because of the impaired kinetochore-microtubule interaction in *Ube2i Zp3-cre+* oocytes, we further analyzed changes in acetylation of histone H4 at lysine 16 (H4K16ac), as this histone modification maintains kinetochore function in somatic cells and possibly also in oocytes (*26, 27*).Control and *Ube2i Zp3-cre+* oocytes were analyzed in GV (Fig. 3D-F), pro-metaphase (Fig. 3G-I), and MI (Fig. 3J-L) by immunofluorescence and quantified by confocal microscopy. The relative amount of H4K16ac showed no difference between genotypes at any stage, indicating that *Ube2i Zp3- cre+* oocytes are more likely to have reduced kinetochore function than control oocytes despite the similarity of H4K16ac immunostaining at the GV, pro-metaphase, or MI stages.

**Fig. 3.**
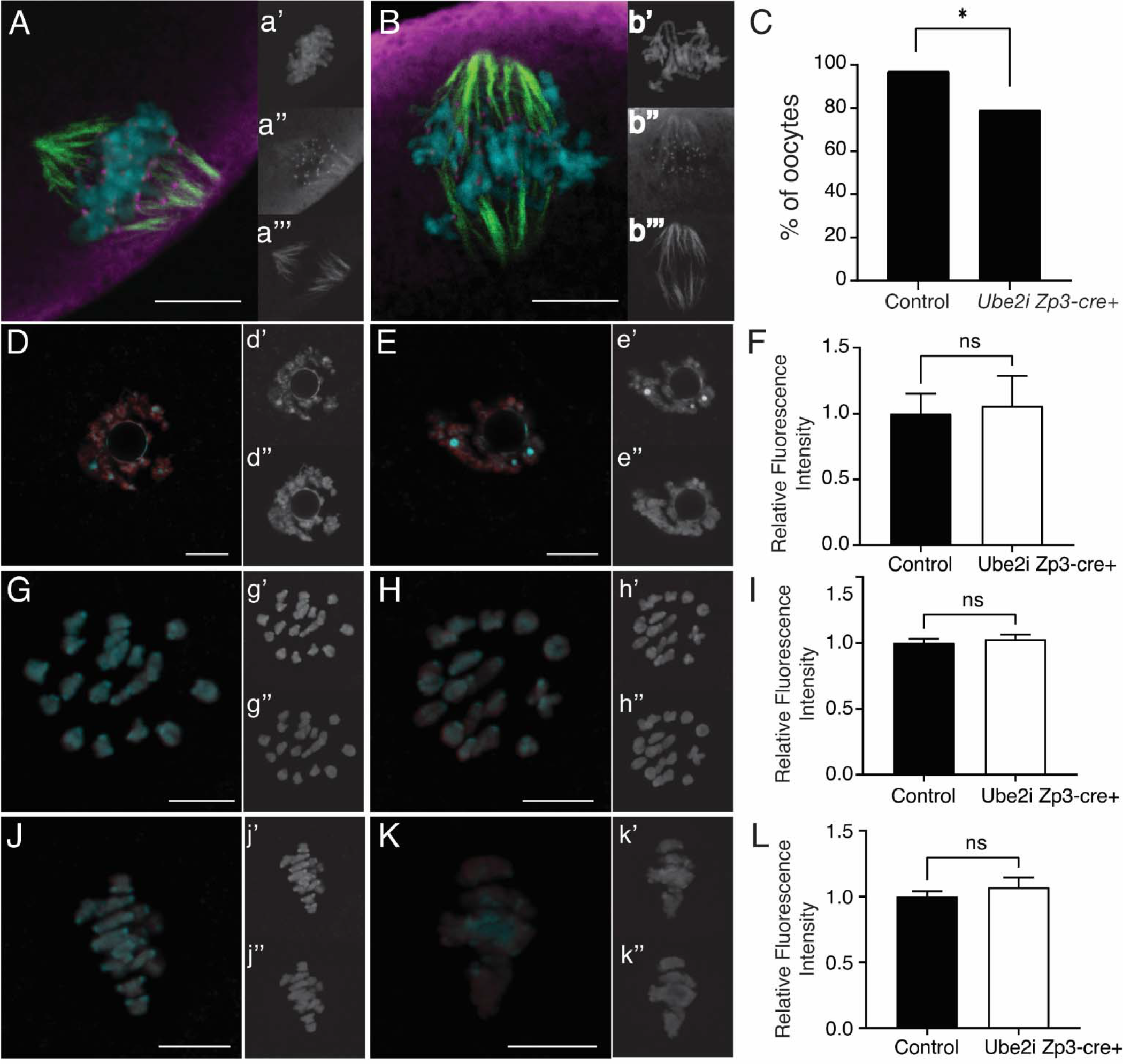
*Ube2i Zp3-cre+* metaphase I oocytes have reduced kinetochore function and normal levels of H4K16ac. Oocytes were collected and cultured for 7 hours, then placed in ice-cold MEM for 10 minutes to depolymerize non-kinetochore microtubules. Oocytes were stained for -tubulin (green), α CREST to mark kinetochores (magenta), and DNA (blue). (A) Merged image of projected representative confocal z-slices of a control cold-treated metaphase I oocyte. (B) Merged image of projected representative confocal z-slices of a *Ube2i Zp3-cre+* cold-treated metaphase I oocyte. Scale bars are 10µm. (C) Quantification of cold-treated oocytes with cold-stable microtubules present. Significantly more control oocytes contained cold-stable microtubules (97.2%) than *Ube2i Zp3-cre+* oocytes (79.2%), indicating reduced kinetochore function in *Ube2i Zp3-cre+* oocytes. (P=0.0333, control n=36, *Ube2i Zp3-cre+* n=24, Fisher’s exact test.) GV oocytes were collected and fixed immediately (GV stage, D, E), cultured to pro-metaphase (G, H), or metaphase I (J, K) then were fixed and stained for H4K16ac (red) and DNA (blue). (D) Merged image of representative z-slice for H4K16ac immunoreactivity (red) in a control (D) and *Ube2i Zp3-cre+* (E) GV oocytes counterstained with Hoechst (blue); unmerged panels are shown in d’ and e’ (Hoechst) and d” and e’’ (H4K16ac). Merged image of representative z- slice for H4K16ac immunoreactivity (red) in a control (G) and *Ube2i Zp3-cre+*(H) pro-metaphase oocyte counterstained with Hoechst (blue); unmerged panels are shown in g’ and h’ (Hoechst) and g” and h’’ (H4K16ac). Merged image of representative z-slice for H4K16ac immunoreactivity (red) in a control (J) and *Ube2i Zp3-cre+* (K) MI oocyte counterstained with Hoechst (blue); unmerged panels are shown in j’ and k’ (Hoechst) and j” and k’’ (H4K16ac). Quantification of positive H4K16ac signal shows no difference in the relative amount of H4K16 at the GV (F) (control n= 35, Ube2i Zp3-cre+ n=29, P>0.05), pro-metaphase (I) (control n= 24, Ube2i Zp3-cre+ n=36, P>0.05), or MI (L) (control n= 30 Ube2i Zp3-cre+ n=25) stages. Scale bars on all images are 10µm. Unpaired t-test.

### *Ube2i Zp3-cre+* GV oocytes fail to compact chromosomes and silence transcription

Prior to meiotic resumption, GV oocytes undergo chromatin remodeling, which changes the configuration of the chromatin around the nucleolus. This change involves the transition from a less compact, non-surrounded nucleolus (“NSN”) chromatin configuration, to a more compact, surrounded nucleolus (“SN”) configuration (*28–30*). We classified oocytes from control and *Ube2i Zp3-cre+* mice as SN - a ring of condensed chromatin surrounding the nucleolus (Fig. 4A, 4C), or NSN - no ring or only a partial ring of condensed chromatin around the nucleolus (Fig. 4B, 4D). Compared to control oocytes, a significantly greater proportion of GV oocytes collected from *Ube2i Zp3-cre+* mice failed to adopt the mature SN configuration and remained in the NSN configuration (Fig. 4E, P < 0.0001). The transition from NSN-to-SN chromatin configuration is associated with the silencing of transcription that precedes the resumption of meiosis (*28*). Due to the failure of *Ube2i Zp3-cre+* oocytes to adopt the SN chromatin configuration, we analyzed transcriptional activity using 5-ethynyl-uridine (5EU) labeling. In control GV oocytes, transcriptionally inactive oocytes showed no fluorescent EU signal (Fig. 5A, 5C), while oocytes that are transcriptionally active have a positive fluorescent signal (Fig. 5B, 5D). *Ube2i Zp3-cre+* oocytes had a significantly higher proportion of oocytes that were transcriptionally active (77%, P<0.0001) compared to control oocytes (28%) (Fig. 5E), suggesting that *Ube2i Zp3-cre+* GV oocytes fail to suppress transcription.

**Fig. 4.**
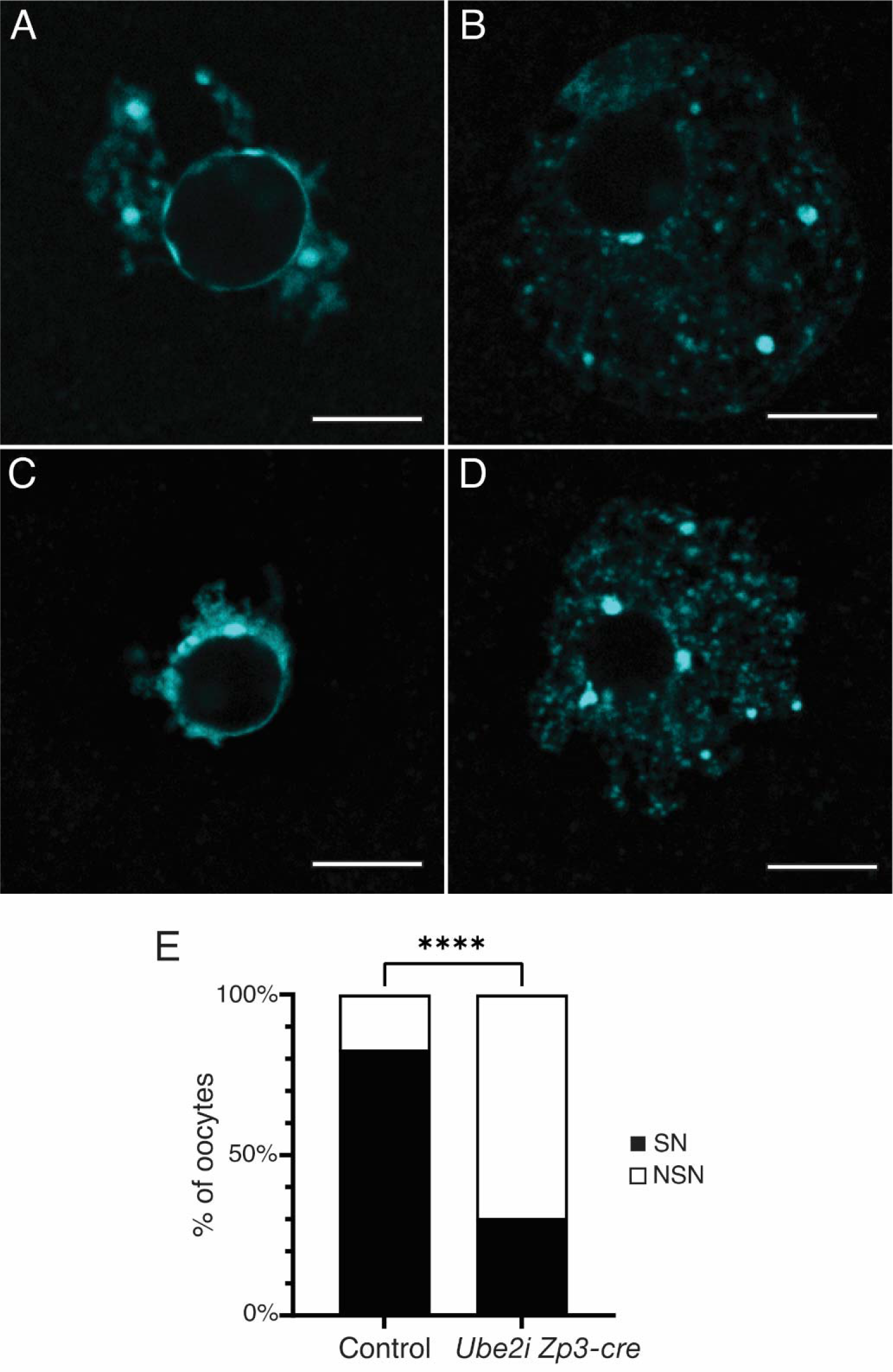
*Ube2i Zp3-cre+* oocytes fail to condense chromatin. GV oocytes collected from control and *Ube2i Zp3-cre+* oocytes were collected, fixed, and stained with Hoechst to analyze DNA configuration. Oocytes with a solid ring of DNA around the nucleolus were classified as surrounded nucleolus (SN) and oocytes lacking or with a partial ring of DNA were classified as non-surrounded nucleolus (NSN). Examples of SN control (A) and SN *Ube2i Zp3-cre+* (C) oocytes. Examples of NSN control (B) and NSN *Ube2i Zp3cre+* (D) oocytes. Representative confocal z-slices, all scale bars are 10µm. (E) Quantification of SN and NSN oocytes for each genotype. (****P < 0.0001, Fisher’s exact test. Control n = 47, *Ube2i Zp3-cre+* n = 46.) SN, surrounded nucleolus; NSN, non-surrounded nucleolus.

**Fig. 5.**
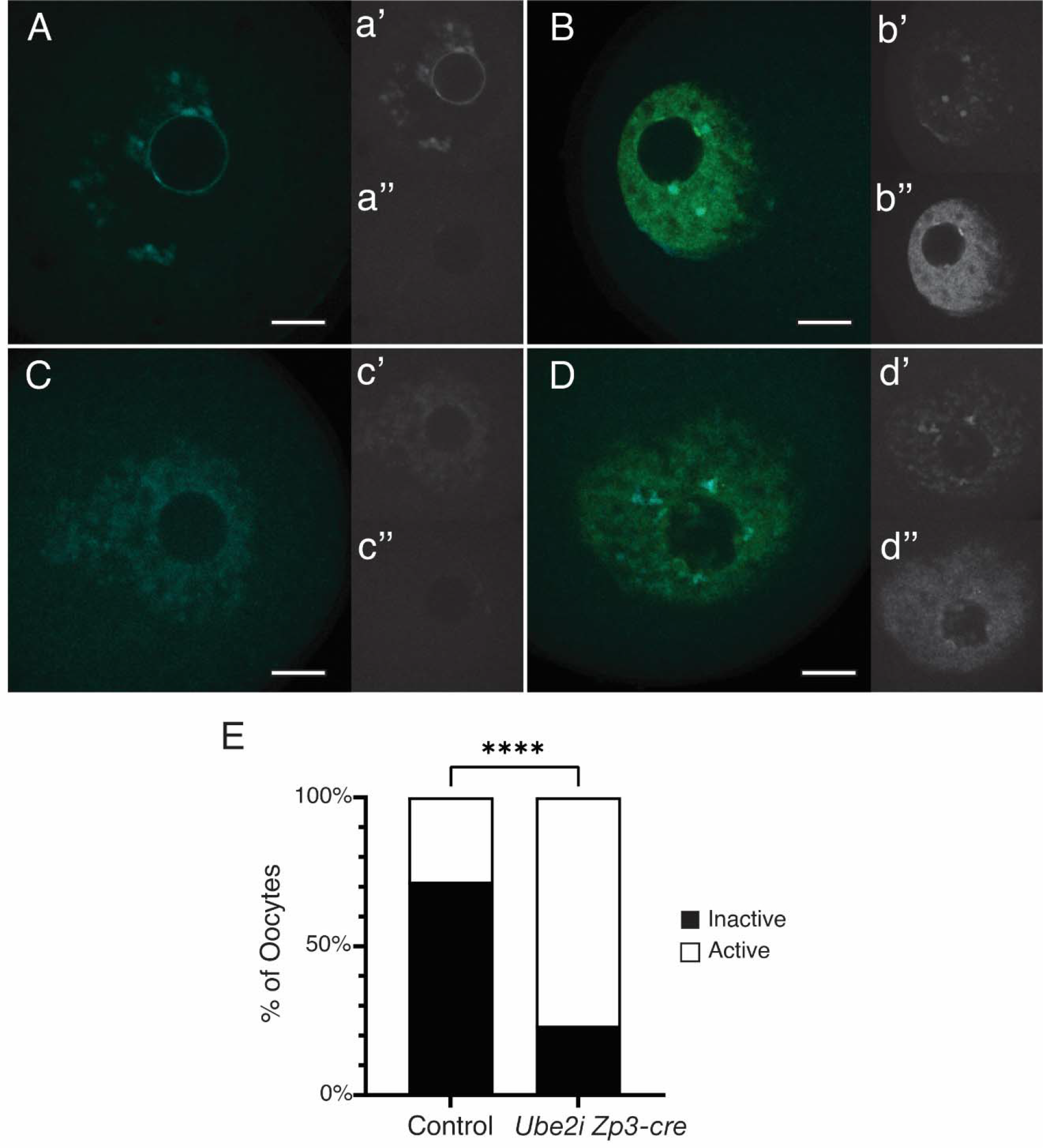
*Ube2i Zp3-cre+* oocytes fail to silence transcription. GV oocytes were held in meiotic arrest with milrinone, and cultured for 2 min in the presence of 5EU to label newly synthesized RNA (positive transcriptional activity), labeled green. DNA was stained with Hoechst. Examples of transcriptionally inactive control (A) and *Ube2i Zp3-cre+* (C). Examples of transcriptionally active control (B) and *Ube2i Zp3-cre+* (D) oocytes are shown for comparison. A significantly higher number of *Ube2i Zp3-cre+* oocytes failed to silence transcription (green). For each image (A-D), the top small insets show DNA (a’, b’, c’, d’), and the bottom small inset shows 5EU signal (a”, b”, c”, d”). Representative confocal z- slices, all scale bars are 10µm. E) Quantification of inactive vs. active transcription for each genotype. (****P < 0.001, Fisher’s exact test. Control n = 46, *Ube2i Zp3-cre+* n = 30.)

Histone modifications such as methylation are associated with transcriptional activity in GV oocytes, and trimethylation of histone H3 at lysine 3 (H3K4) and lysine 9 (H3K9) in mouse oocytes, H3K4me3 and H3K9me3 are markers of transcriptional repression (*31*). Because of the active transcription observed in *Ube2i Zp3-cre+* oocytes (Fig. 5), we analyzed H3K4me3 and H3K9me3 levels in GV oocytes by immunofluorescence (Fig. 6). Compared to control oocytes, *Ube2i Zp3-cre+*oocytes had significantly reduced levels of both H3K4me3 (34%, P<0.0001, Fig. 6A-C) and H3K9me3 (27%, P<0.0001, Fig. 6D-F). Repressive histone marks such as H3K4me3 and H3K9me3 are associated with histone H3.3 in mouse oocytes (*31*). To determine if the reduction of H3K4me3 in *Ube2i Zp3-cre+* oocytes is due to the failure of H3.3 incorporation into nucleosomes, GV oocytes were analyzed for histone H3.3 by quantitative whole-mount immunofluorescent analysis, whichprovides a positive signal for histone H3.3 incorporated into nucleosomes, as well as free histone H3.3 in the nucleus. Compared to control oocytes, *Ube2i Zp3-cre+* oocytes had a small (15.1%) but statistically significant decrease in the amount of free histone H3.3 in the nucleus compared to control oocytes (Fig. S2).

**Fig. 6.**
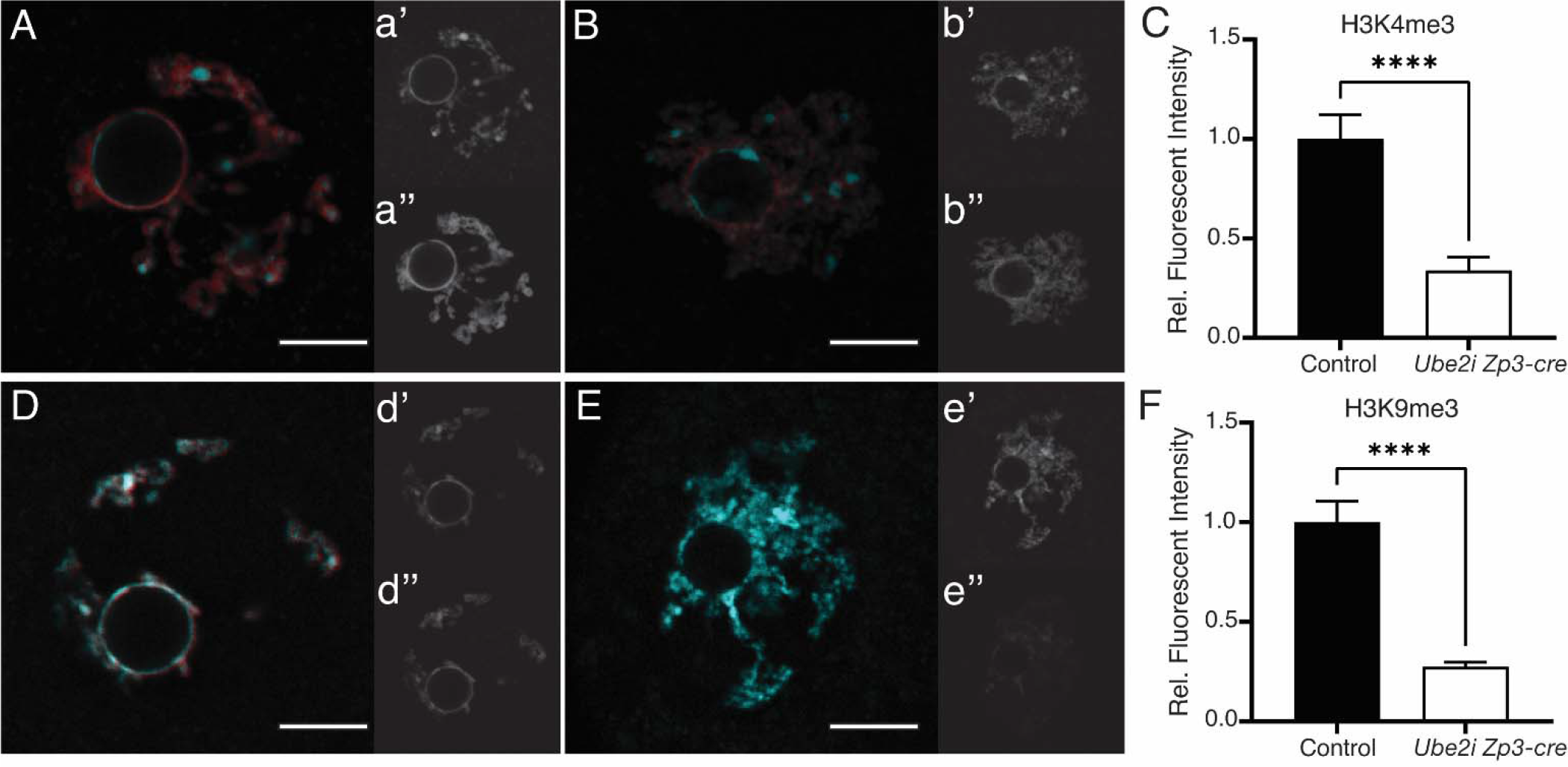
Repressive histone modifications H3K4me3 and H3K9me3 are significantly decreased in *Ube2i Zp3-cre+* oocytes. Control and *Ube2i Zp3-cre+* GV oocytes were collected and immunostained for H3K4me3 (top row) or H3K9me3 (bottom row) and counterstained for DNA (Hoechst) (blue). (A) Merged image of representative z-slice for H3K4me3 immunoreactivity (red) in a control oocyte counterstained with Hoechst (blue); unmerged panels are shown in a’ (Hoechst) and a” (H3K4me). (B) Merged image of representative z-slice for H3K4me3 (red) immunoreactivity in a *Ube2i Zp3+* oocyte counterstained with Hoechst (blue); unmerged panels are shown in b’ (Hoechst) and b” (H3K4me). (D) Merged image of representative z-slice for H3K9me3 immunoreactivity (red) in a control oocyte counterstained with Hoechst (blue); unmerged panels are shown in d’ (Hoechst) and d” (H3K9me). (E) Merged image of representative z-slice for H3K9me3 (red) immunoreactivity in a *Ube2i Zp3+* oocyte counterstained with Hoechst (blue); unmerged panels are shown in e’ (Hoechst) and e” (H3K9me). (C,F) Quantification of positive H3K4me3 (C) or H3K9me3 (F) signal shows that the abundance of both modifications is significantly reduced in *Ube2i Zp3-cre+* oocytes. Representative confocal z-slices, all scale bars are 10µm. (H3K4me3 staining, ****P < 0.0001, unpaired t-test. Control n = 31, *Ube2i Zp3-cre+* n = 30.) (H3K9me3 staining ****P < 0.0001, unpaired t-test. Control n = 31, *Ube2i Zp3-cre+* n = 28.)

### *Ube2i Zp3-cre+* GV oocytes have a significantly altered transcriptome

Because *Ube2i Zp3-cre+* GV oocytes fail to silence transcription, we next determined if the transcriptome was alterend between control and *Ube2i Zp3-cre+* oocytes. RNA-sequencing of individual GV stage oocytes was performed on control and *Ube2i Zp3-cre+* GV oocytes (n=17 per genotype). Principal component analysis (PCA) identified two groups with 70% variance in PC1, which readily separates the samples by genotype; 15% variance is explained by PC2, which showed a wider sample spread for *Ube2i Zp3-cre+* oocytes than control (Fig 7A). We identified 7,604 differentially expressed genes (DEG) in *Ube2i Zp3-cre+* GV oocytes compared to control oocytes using a cutoff of log2 (Fold Change) and P<0.05 (Fig. 7B). Of these DEG, more upregulated genes (6, 309) were ≥ found compared to downregulated genes (1, 295), supporting the hypothesis that *Ube2i Zp3-cre+* GV oocytes remain transcriptionally active.

**Fig. 7.**
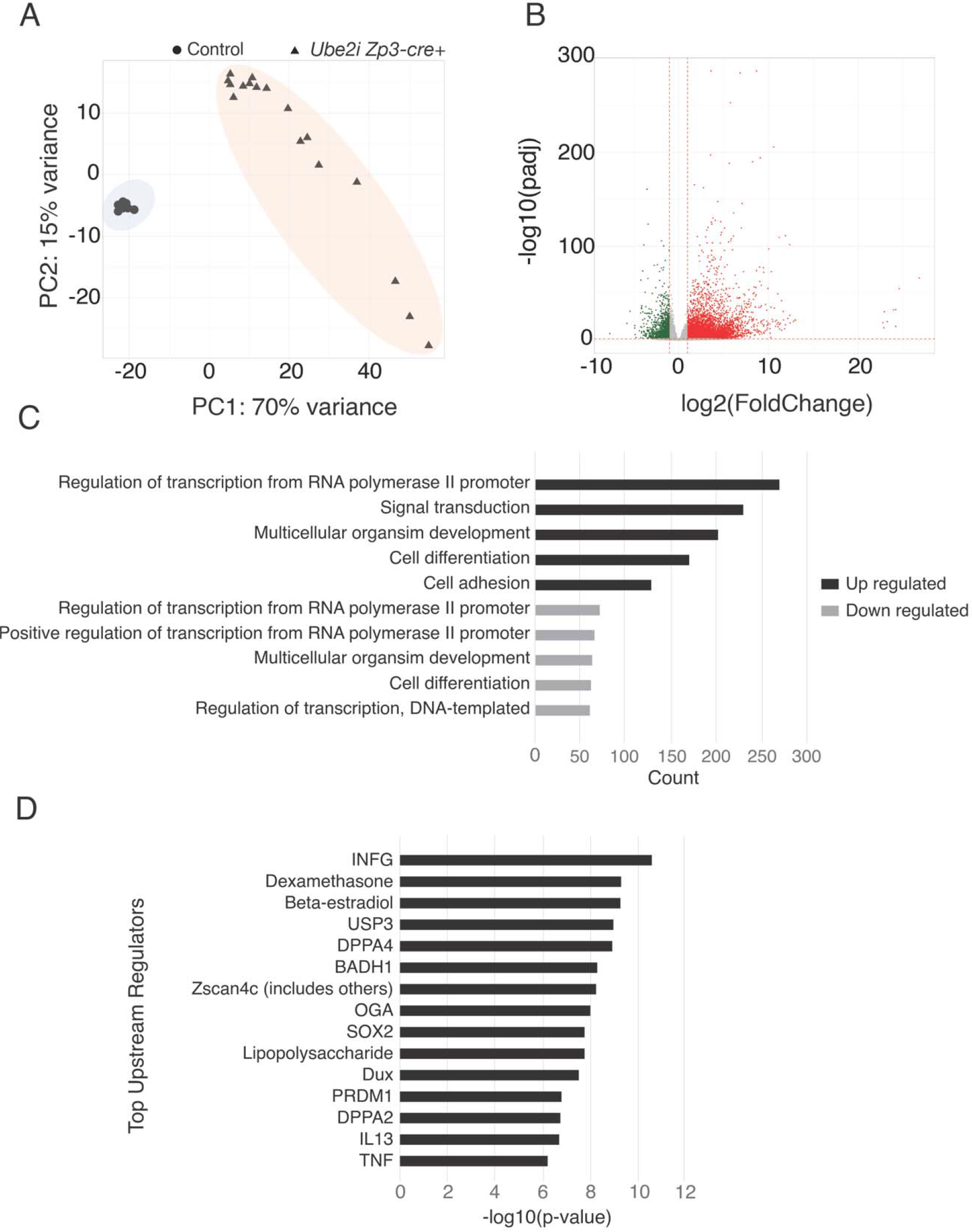
*Ube2i Zp3-cre+* oocytes have a significantly altered transcriptome. (A) Principal component analysis plot of control and *Ube2i Zp3-cre+* oocytes. (B) Volcano plot showing differentially regulated genes (DEG) in control and *Ube2i Zp3-cre+* GV oocytes from RNA-sequencing of individual oocytes. C) Gene ontology (GO) analysis of DEG with a 2-fold increase or a 2-fold decrease, p-value < 0.05. D) Top upstream regulators determined from Ingenuity Pathway Analysis (IPA) of RNA-sequencing data.

To investigate biological functions associated with the DEG, gene ontology (GO) analysis was performed for differentially upregulated and downregulated DEG (*32*). Separate analysis of both upregulated and downregulated genes returned the same top GO term: *regulation of transcription from RNA polymerase II promoter*. Differentially upregulated genes also returned the GO terms *signal transduction*, *multicellular organism development*, *cell differentiation*, and *cell adhesion*, while downregulated genes returned the GO terms *positive regulation of transcription from RNA polymerase II promoter*, *multicellular organism development*, *cell differentiation*, and *DNA-templated regulation of transcription* (Fig. 7C).

Key oocyte-expressed genes were differentially regulated in *Ube2i Zp3-cre+* GV oocytes (Fig. 8A). Genes downregulated in *Ube2i Zp3-cre+* GV oocytes included *Bmp15*, *Gdf9*, *H1oo*, *Hormad1*, *Mater*, *Npm2*, *Oosp1*, *Plk1*, *Rfpl4*, *Tgfb2*, *Msy2*, *Zar1*, and *Zp1-3,* while upregulated genes include *Blimp1, Pou5f1/Oct4* and *Trim28*. Of note, both *Bmp15* and *Gdf9* are also significantly downregulated in *Ube2i Gdf9-icre_+_* ovaries (*18*)and may contribute to the less well-developed cumulus cell layer observed in antral follicles of *Ube2i Zp3-cre+* ovaries (Fig. S1F). The oocyte-specific transcription factor, *Nobox,* was significantly upregulated in *Ube2i Zp3-cre+*GV oocytes as it is in *Ube2i Gdf9-icre_+_* ovaries (Fig. 8B), however it was the only oocyte-specific transcription factor found to be differentially regulated. Histone lysine methyltransferases and demethylases known to regulated trimethylation of H3K4 and H3K9 were also analyzed in the DEG list. The methyltransferases that were upregulated in *Ube2i Zp3-cre+* oocytes included *Kmt2e*, *Setd1b*, *Setdb1*, and *Setdb2*, and downregulated were *Setd1a*, *Smyd2*, and *Suv39h2* (Fig. S3A). The demethylases that were upregulated include *Kdm4b*, *Kdm4c*, *Kdm4dl*, *Kdm5a,* and *Kdm5b*, and downregulated were *Kdm2b* and *Kdm7a* (Fig. S3B).

**Fig. 8.**
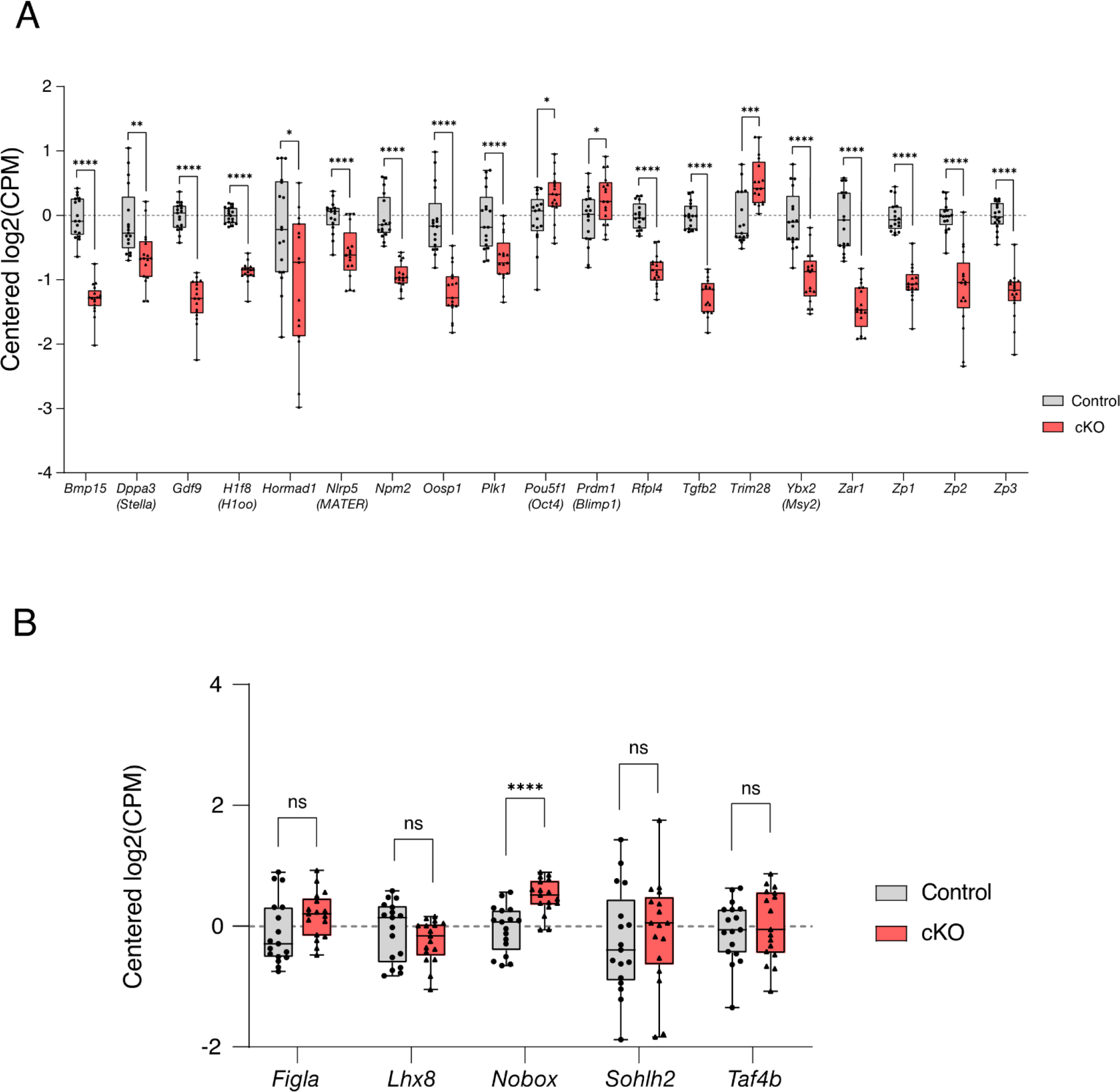
Selected differentially expressed genes (DEG) in *Ube2i Zp3-cre+* GV oocytes. A) Selected genes known to be important in oocyte development and fertility were found to have significantly altered expression levels in *Ube2i Zp3-cre+* GV oocytes compared to control GV oocytes. B) The only oocyte-specific transcription factor found to be differentially regulated is *Nobox*, which has higher expression in in *Ube2i Zp3-cre+* GV oocytes.

Major chromatin remodeling is not required for global transcriptional silencing in GV oocytes (*33*). However, to verify that transcriptome changes were not due to persistence of the NSN configuration of mutant GV oocytes, the DEG list from *Ube2i Zp3-cre+* oocytes was comparedgene expression profiles previously reported to be associated with SN and NSN oocytes (*28*). The genes downregulated in wild type SN compared to NSN is only 2% (152/7604) of the DEG in *Ube2i Zp3-cre+* oocytes. Similarly, genes upregulated in wild oocytes from NSN to SN comprise a mere 3% (226/7604) of the *Ube2i Zp3-cre+* oocytes changes (Fig. S5). Thus, the predominance of NSN in *Ube2i Zp3-cre+* oocytes is unlikely to explain the large transcriptome changes in the mutant oocytes. (Fig. S4).

### Loss of *Ube2i* in oocytes results in the premature activation of the zygotic transcription program

To further analyze the differences in gene expression, Ingenuity Pathway Analysis (IPA) of the DEG was used to identify top upstream regulators. This analysis revealed that several of the top upstream regulators were associated with zygotic genome activation (ZGA), including *Dppa2, Dppa4*, *Dux*, *Sox2*, and *Zscan4* (Fig. 7D) (*34*). While *Dppa4* expression was significantly decreased in *Ube2i Zp3-cre+*oocytes, *Zscan4* (including *Zscan4a-f*), *Sox2*, and *Duxf3* were significantly increased, and *Dppa2* was unchanged between genotypes. The ZGA-associated genes, *Duxf3* and *Zscan4,* were validated in independent samples of control and *Ube2i Zp3-cre+* GV oocytes. By qRT-PCR, we confirmed that these genes were upregulated in *Ube2i Zp3-cre+* oocytes compared to control oocytes (Fig. 9A, 9B). Other ZGA associated genes such as *Eif1a*, an embryonic transcription factor, and *Tmem92*, a downstream target of ZSCAN4, had significantly higher expression in both the list of DEG from the RNA-seq dataset as well with qRT-PCR validation (Fig. 9C, 9D). Finally, to further test activation of the zygotic genome, *MuERV-L*, one of the earliest transcribed genes in mouse one-cell embryos, was analyzed by qRT-PCR and found to be significantly upregulated in *Ube2i Zp3-cre+* oocytes (4,014% compared to control, P < 0.0001, Fig. 9E) (*35*). These data demonstrate that UBE2I is essential to silence transcription in GV oocytes and to prevent the premature activation of the zygotic transcription program.

**Fig. 9.**
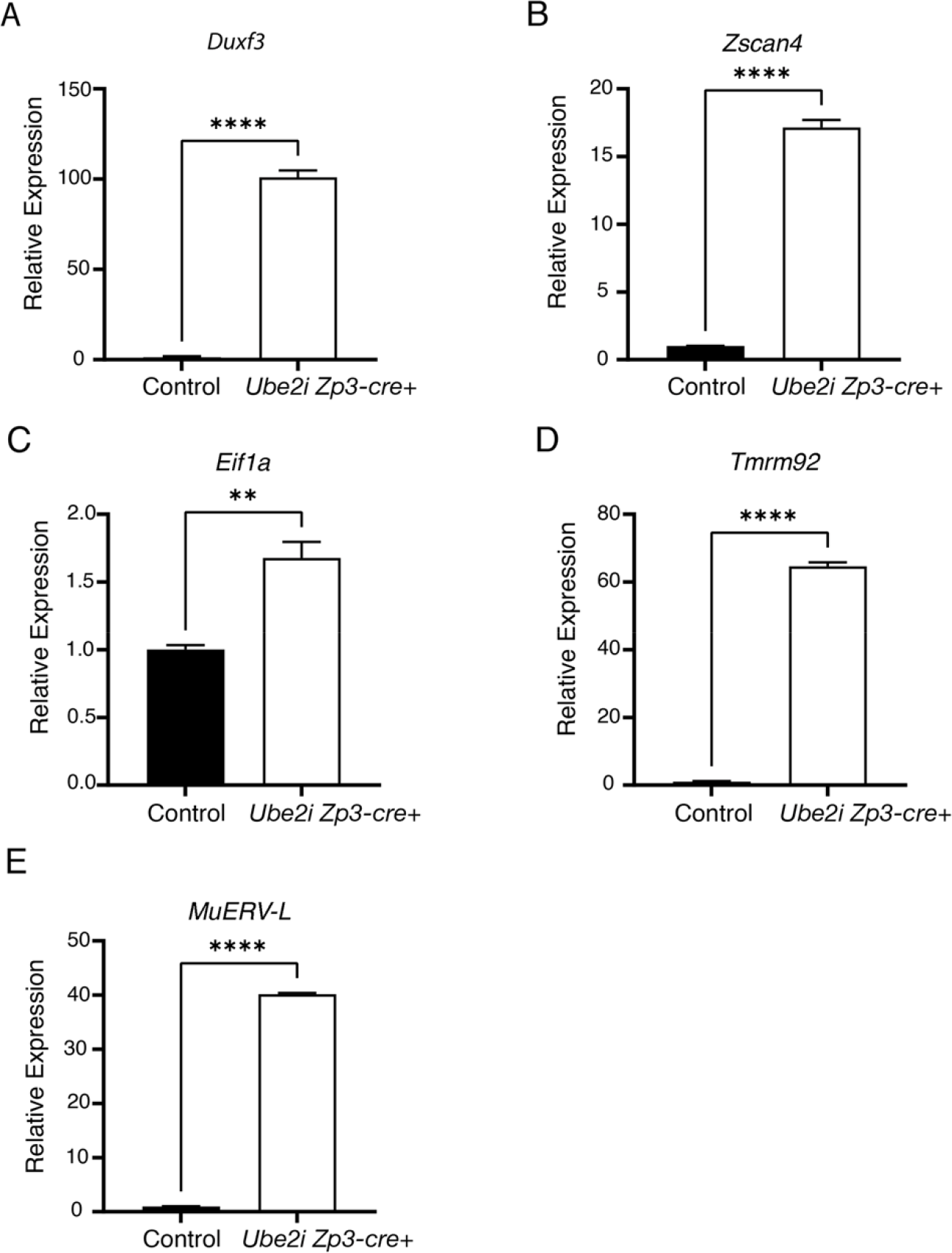
GV *Ube2i Zp3-cre+* oocytes express multiple genes associated with the activation of the zygotic genome. cDNA was generated from RNA isolated from GV oocytes from control or *Ube2i Zp3- cre+* females to validate the expression of selected genes from the RNA-sequencing data. *Duxf3* (A), *Zscan4* (B), *Eif1a* (C), and *Tmem92* (D) expression was verified to be significantly upregulated in *Ube2i Zp3-cre+* oocytes. E) *MuERV-L*, one of the earliest expressed genes in the one-cell mouse embryo, is significantly upregulated in *Ube2i Zp3-cre+* oocytes (**P = 0.0053, ****P < 0.0001, unpaired t-test).

## Discussion

The scope of regulation provided by SUMOylation in mammalian oocyte biology is not well known, particularly during oocyte growth within ovarian follicles. This time period is critical for establishing key hallmarks of oocyte development, including acquisition of meiotic and developmental competence. In addition, during folliculogenesis, there are major changes in chromatin organization (NSN to SN) related to increased meiotic and developmental competence (*29, 36*). In addition, the SN configuration is associated with global suppression of transcription (*3, 37, 38*). While SUMOylation is associated with transcriptional repression in general (*39, 40*), a role for SUMOylation in global transcriptional repression in oocytes has not been previously described. Using a novel mouse strain developed to delete *Ube2i* selectively in growing oocytes, we demonstrate that UBE2I is essential in chromatin remodeling, meiotic progression, and transcriptional regulation.

Oocyte development is an extended process, beginning in the embryo, continuing during ovarian folliculogenesis, and not completing until fertilization. We previously developed an oocyte-specific knockout of *Ube2i* using *Gdf9-icre* and discovered that *Ube2i* was required for stability of the ovarian reserve, regulation of the oocyte-specific transcriptional program, and resumption of meiosis (*18*).

However, the premature loss of ovarian follicles in early adulthood hampered our ability to study *Ube2i* in later oocyte developmental stages. Furthermore, it was unclear if accelerated depletion of the ovarian reserve was due to direct effects of *Ube2i* in oocytes of primordial follicles, or indirectly due to loss of the growing follicle populations, which are known to produce factors, such as anti-Müllerian hormone that regulate dynamics of the primordial follicle (*41, 42*). Therefore, to address this gap, we utilized the well-characterized *Zp3-cre* deleter strain for oocytes of growing follicles already recruited from the primordial follicle pool (*24*). Even though both models are display female sterility with oocytes that arrest at MI, in contrast to *Ube2i Gdf9-icre*+ ovaries (*18*), *Ube2i Zp3-cre+* ovaries show a stable primordial follicle pool with minimal effects on follicle dynamics. Comparing these two models, it is clear that *Ube2i* has essential roles in oocytes of primordial follicles and in oocytes of growing follicles. Furthermore, a comparison of these models highlights that oocytes in the non-growing primordial follicles are far from ‘quiescent’, and defects accumulated in oocytes prior to being recruited for folliculogenesis leads to poor quality oocytes with a decreased levels of meiotic resumption (*18*). For a further comparison between *Ube2i Gdf9-icre*+ and *Ube2i Zp3-cre+*models, see supplementary table S1.

In contrast to *Ube2i Gdf9-icre*+ MI spindles (*18*), *Ube2i Zp3-cre+* MI spindles are abnormal, similar to defects identified when SUMOylation is acutely inhibited *in vitro* (*21, 43, 44*). *Ube2i Zp3- cre+* spindles are both shorter and narrower than control spindles, in addition to showing poor alignment of chromosomes on the metaphase plate. One contributor to the poor spindle formation in *Ube2i Zp3- cre+* maybe be decreased expression and/or activity of polo-like kinase 1 (*Plk1*). a SUMO target and overexpression of PLK1 SUMOylation site mutants in oocytes causes increases in the number of abnormal spindles (*21, 45, 46*). In oocytes, PLK1 coordinates key events during meiotic resumption and progression, including regulation of chromosome architecture and spindle assembly. Oocyte-specific deletion of *Plk1* in mice causes female sterility and MI arrest and *Plk1* conditional knockout (cKO) oocytes contain abnormally shaped (i.e., reduced width and length) MI spindles (*47*). Both *Ube2i Gdf9- icre+* ovaries and *Ube2i Zp3-cre+* oocytes show downregulation of *Plk1*. Thus, SUMOylation of *Plk1* to regulate spindle assembly is well supported, while remaining studies are needed to clarify mechanisms underlying additional meiotic defects in *Ube2i Zp3-cre* that arise independent of PLK1 levels or its modification by SUMOylation. *Ube2i Zp3-cre+* spindles also show reduced ability to form proper microtubule-kinetochore attachment as measured by the presence or absence of cold-stable microtubules at MI. Kinetochore function in mouse oocytes has been reported to be regulated by H4K16 acetylation mediated by HDAC2, as increased levels of H4K16ac in *Hdac2^-/-^* or *Hdac1^-/+^ Hdac2^-/-^* MI oocytes reduces kinetochore-microtubule attachments (*26*)Although no change in H4K16ac levels were observed at the GV, pro-metaphase, or metaphase I stages, *Ube2i Zp3-cre+* metaphase I oocytes were significantly more likely to lack cold-stable microtubules than control oocytes. The normal levels of H4K16ac indicates that HDAC2 is functioning in *Ube2i Zp3-cre+* oocytes, suggesting that the reduction in cold-stable microtubules observed at metaphase I is the result of an alternative kinetochore-regulating mechanism, perhaps through PLK1 (*48–50*).

Another key defect in *Ube2i Zp3-cre+*GV oocytes is the failure to transition from the NSN to SN chromatin configuration. In antral follicles, a significant proportion of fully grown oocytes are in the SN configuration, with the remainder in NSN (*51*). While wild type NSN oocytes progress normally through meiosis and can be fertilized, they subsequently arrest at the 2-cell stage, resulting in no viable offspring (*36*). In contrast, the majority of *Ube2i Zp3-cre+* GV oocytes show a two-fold reductions in meiotic resumption despite the failure to adopt the SN configuration. Of the oocyte-expressed genes that are downregulated in *Ube2i Zp3-cre+,* several have knockout mouse models that also show an increased percentage of GV oocytes in the NSN configuration, including knockouts of the maternal effect genes, *Npm2* and *Mater,* both of which arrest at the 2-cell embryo stage (*52–54*). The maternal effect gene, *Zar1* for which the knockout mouse model arrests at the one-cell embryo stage (*55*), is also downregulated in *Ube2i Zp3-cre+* GV oocytes. Interestingly, *Zar1* and *Mater* are degraded from GV to MII (*56*), with 90% of maternal effect genes are degraded by ZGA (*57, 58*). The decreased levels of maternal effect transcripts in spite of the maintenance of active transcription in *Ube2i Zp3-cre+* may suggest that RNA stability or storage may be affected. The RNA-binding protein, MSY2, is significantly downregulated ∼2-fold, and *Msy2^-/-^* GV oocytes have a 25% reduction in poly(A)-containing RNA (*59*). Why *Msy2* is downregulated in *Ube2i Zp3-cre+* remains to be determined. RNA binding proteins (RBP) are also targets of SUMOylation or contain SUMO-interacting motifs (*60, 61*), however, RBP SUMO-targets in oocyte biology remains to be determined.

The NSN to SN chromatin configuration switch of fully grown oocyte changes is associated with transcriptional silencing (*28, 37*), though these processes can be uncoupled, suggesting they have different regulatory mechanism (*33*). The mechanisms underlying the transcriptional silencing are not well understood (*62*). *Ube2i Zp3-cre+* GV oocytes fail to silence transcription with +6000 genes differentially upregulated, although some gene classes, such as the maternal effect genes are downregulated. The ongoing transcription is not simply a result of a change in the ratio of NSN to SN because there is little overlap between transcriptome gene sets comparing SN to NSN oocytes and our single oocyte RNA-sequencing results (*28*). Previously, overexpression of *Ube2i* in GV mouse oocytes initially indicated that UBE2i stimulated transcription (*8*); however, the model presented here demonstrates that UBE2I is required to silence transcription. This latter model is similar to studies in embryonic stem cells, which show that SUMOylation functions in heterochromatin to maintain proper genome-wide H3K9me3 by triggering repressive complexes that contain SUMOylated PRC1.6 complex and TRIM28(KAP)/SETDB1. In both *Ube2i Gdf9icre+*and *Ube2i Zp3cre+*models, PRDM1 (also called BLIMP1) appears as a top upstream regulator of the DEG lists. PRDM1 is a is zinc finder SET domain protein that acts as a transcriptional repressor and controls cell fate decisions (*63*) In *Ube2i Zp3cre+ GV oocytes, Prdm1* is slightly but significantly upregulated 1.2-fold relative to controls.

PRDM1 is SUMOylated in mouse B-cells, and its SUMO-resistant mutant of PRDM1 causes a failure to silence transcription and impairs differentiation (*64*). *Prdm1^-/-^* mice die embryonically with impaired primordial germ cell development (*65*), but it has not been characterized in oocyte development.

Transcriptional silencing in wild type GV oocytes is associated with the accumulation of H3K4 and H3K9 trimethylation. *Ube2i Zp3-cre+* GV oocytes show loss of these repressive marks as well as failure to silence transcription, implicating SUMOylation in the establishment of trimethylated H3K4 and H3K9 and transcriptional silencing. Interestingly, decreased levels of H3K4me3 and H3K9me3 is also associated with expression of ZGA associated genes such as *Zscan4* (*31*). Indeed, active removal of H3K4me3 by the lysine demethylases, KDM5A and KDM5B, is required for ZGA, and reduction of these marks results in premature activation of the ZGA in GV oocytes (*66*). Informatic analysis of *Ube2i Zp3-cre+* GV oocyte DEG revealed ZGA associated genes as top upstream regulators, suggesting the increased transcription in *Ube2i Zp3-cre+* is, in part, due to premature activation of the zygotic genome. In support of a regulatory role for SUMOylation in ZGA, a prior study demonstrated that overexpression of the SUMO E3 ligase, PIASy, in two-cell embryos increased H3K9me3, disrupted the ZGA as indicated by reduced expression of MERLV transcripts (one of the earliest transcribed genes in the minor ZGA), and arrests cells at the two-cell stage (*67*). Other studies have showed that SUMOylation regulates KDM5B incultured cells and prevents it from binding to gene promoter regions, while SUMO- deficient KDM5B occupies target gene promotors leading to demethylation of H3K4me3 and gene silencing (*68*). These studies, in tandem with the *Ube2i Zp3-cre+* phenotype of maintenance of active transcription alongside premature ZGA, indicates that SUMOylation is required to silence transcription in fully grown oocytes prior to meiotic resumption and that the lack of SUMOylation results in a permissive environment for ZGA. Additionally, ZGA activation in *Ube2i Zp3cre+ GV* included upregulation of *Zscan4*, *Eif1ad15(Gm5039)*, *Duxf3*, *Sox2*, and *MuErv-L*, and a similar upregulation of *Dux*, *Zscan4*, and *Eif1a-like* genes is noted when *Ube2i* is knocked down in embryonic stem cells, alongside a decrease in global H3K9me, the result of which converts cells to a more pluripotent, 2-cell embryo like state (*13*). Taken together with the decrease in histone H3K4me3 and H3K9me3 in *Ube2i Zp3-cre+* oocytes and the apparent premature activation of the zygotic genome in *Ube2i* GV oocytes, further work to determine if any of these proteins are regulated by SUMOylation in mouse oocytes will be crucial.

Major regulatory pathways controlling transcriptional silencing in fully grown oocytes as well as triggers for maternal RNA degradation and ZGA are not fully defined. The continued transcriptional activity and upregulation of ZGA genes in GV oocytes lacking *Ube2i* indicates a critical role for UBE2I in all three processes: silencing transcription prior to meiotic resumption, regulation of maternal effect gene transcripts, and the resumption of transcription in early embryos during the maternal-to-zygotic transition. Our data from the *Ube2i Zp3-cre* model suggest that SUMOylation is a key pathway for silencing oocyte transcription at the end of folliculogenesis, while de-SUMOylation may be required for degradation of maternal transcripts and activation of transcription in the early embryo. Thus, SUMOylation may safeguard oocyte ‘fate’ until it is necessary to activate the zygotic genome, similar to its role in other cellular contexts (*13*). Perhaps, in hindsight, it is not surprising that posttranslational modifications such as SUMOylation will turn out to be key determinates of the developmental time frame spanning meiotic resumption, meiotic maturation, fertilization, and preimplantation embryo development. Determining direct SUMO targets, including which histone-methyltransferases and demethylases are modified by SUMOylation and how this affects their function will further uncover roles for global SUMOylation at the key developmental stages required for female fertility.

## Materials & Methods

### Experimental mice

Experimental procedures utilizing mice were performed in compliance with the National Institutes of Health Guide for the Care and Use of Laboratory Animals, and Institutional Animal Care and Use Committee-approved animal protocols at Baylor College of Medicine (AN-4762). The *Ube2i^loxP/loxP^* mice were previously generated and provided by Sean Hartig (*69*). *Ube2i^loxP/loxP^* females were mated to *Zp3-cre+* males to generate *Ube2i^loxP/+^ Zp3-cre+* males, which were then crossed to *Ube2i^loxP/loxP^* females to generate *Ube2i^loxP/loxP^ Zp3-cre+* females. Generation and characterization of *Zp3-cre* mice has been previously characterized (*24*). Genomic DNA from ear punches was used for genotyping as described (*24*). *Zp3-cre* negative female littermates (*Ube2i^loxP/loxP^*) were used as controls in all experiments to minimize differences in genetic backgrounds. All mice were crossed to and maintained on a C57BL/6J/129S7/SvEvBrd genetic background, similar to our previous study (*18*).

*Ube2i^loxP/loxP^* were also crossed to *Gdf9-icre*, and the phenotype validated against the prior study (*18*). Genotyping primers are listed in table S2.

### Fertility analysis

Sexually mature (six-week-old) *Ube2i Zp3-cre+* females or control littermates were pair-housed with a sexually mature (8-wk old) wild-type male of known fertility and continuously mated for six months. Pregnancy and the presence of newborn pups was monitored daily with any resulting litters weaned at 3-weeks of age. The number of pups and litters were recorded for three mice per genotype.

### Tissue collection, histological analysis, and follicle quantification

Mice were anesthetized using isoflurane (Abbott Laboratories) inhalation and euthanized by cervical dislocation. Ovaries were collected and fixed in 10% neutral buffered formalin (Electron Microscopy Services) overnight, followed by standard paraffin processing and embedding at the Human Tissue Acquisition and Pathology Core at Baylor College of Medicine. Six ovaries of each genotype were serially sectioned (5µm sections) and stained with periodic acid (Sigma Aldrich) and Schiff’s reagent (Sigma Aldrich) for analysis. Brightfield images were obtained using Axioscope 2 plus and Zeiss Zen Software v2.3. For morphometric quantification, follicles were manually counted on every fifth section (*70*).To avoid overcounting, only follicles with a visible oocyte nucleus were counted. For morphometric quantification, follicles were manually counted on every fifth section (*70*).To avoid overcounting, only follicles with a visible oocyte nucleus were counted. For follicles smaller than the antral stage, a correction factor was applied Corpora lutea and antral follicles were counted in total by following each through serial images. Follicle classification was based on previous studies into primordial (type 2), primary (types 3a and 3b), secondary (type 4), preantral (types 5a and 5b) and antral (types 6-8) (*71*).Follicle classification was based on previous studies into primordial (type 2), primary (types 3a and 3b), secondary (type 4), preantral (types 5a and 5b) and antral (types 6-8) (*71*).

### Hormone Analysis

Blood was collected from isoflurane-anesthetized six-month-old mice by cardiac puncture, and serum was separated by centrifugation in microtainer collection tubes (Becton, Dickinson and Company) and frozen at -20° until assayed. Mouse FSH and E2 levels were analyzed by ELISA at the University of Virginia Ligand Core Facility (Specialized Cooperative Centers Program in Reproductive Research NICHD/NIH U54-HD28934). Assay method information is available online (https://med.virginia.edu/research-in-reproduction/ligand-assay-analysis-core/assay-methods/). Data were log transformed before statistical analysis by Welch’s t-test (FSH) or unpaired t-test (E2).

### Oocyte collection, culture, and fixation

For analysis of oocytes following ovulation four- to six-week-old female mice were injected with 5IU PMSG (ProSpec Bio), and 46 hours later injected with 5 IU hCG (Pregnyl; Merck Pharmaceuticals). Mice were euthanized 18 hours after injection with hCG and oocytes were harvested from the oviduct ampulla and fixed in 3.7% PFA (Electron Microscopy Sciences) in BSA-PBS (0.1% BSA (Sigma Aldrich) in PBS) for 15-20 minutes then rinsed through five drops of BSA-PBS to prevent over fixation. Oocytes were stored in BSA-PBS at 4°C overnight, or directly used for immunofluorescent staining.

For collection of GVs, four- to six-week-old female mice were injected with 5IU PMSG and ovaries were excised after 44-46 hours into M2 media supplemented with 2.5µM milrinone (Sigma Aldrich) to maintain meiotic arrest. Follicles were ruptured via puncturing with a 26G needle to release GV oocytes into the media. Oocytes were collected and denuded of cumulus cells via mechanical pipetting. After denuding, oocytes were rinsed through 10 drops of CZB media supplemented with 2mM L-glutamine (Thermo Fisher Scientific) to wash out milrinone and cultured in a fresh drop of medium covered with mineral oil (Sigma Aldrich) at 37°C and 5% CO2. Oocytes that had not undergone GVBD were removed from the culture after 2.5 hours, and oocytes were cultured for five hours for pro-metaphase I, and 7.5-8 hours for metaphase I or for monitoring meiotic progression. After culturing to the desired stage, oocytes were fixed in 3.7% PFA in BSA-PBS for 15-20 minutes then rinsed through five drops of BSA-PBS to prevent over fixation. Oocytes were stored in BSA-PBS at 4°C overnight, or directly used for immunofluorescent staining.

### Chromosome Spreads

Oocytes were collected and cultured to prometaphase (5hr) or metaphase I (8.5 hr) as described above, followed by the removal of the zona pellucida (ZP) by transferring the oocytes through up to 10 drops of acid Tyrode’s solution (Sigma Aldrich) under a brightfield microscope until the ZP was visibly removed. After removal the ZP, oocytes were rinsed through 10 drops of M2 media and returned to the incubator to recover for 30 minutes. 50 µL of chromosome spreading solution (0.64% PFA, 3mM dithiothreitol (Thermo Scientific), 0.16% triton X100 (Thermo Fisher Scientific)), was applied to the wells of PTFE printed slides (VWR), and three to five oocytes were transferred to the well. Lysing of the oocytes was confirmed by visual monitoring using a brightfield microscope. The slides were allowed to dry at room temperature and stored at -20°C until use.

### Immunofluorescent staining

After fixation, oocytes were permeabilized in 0.1% Triton X100 in BSA-PBS for 20 minutes, then rinsed through five drops of BSA-PBS. Blocking was performed in 1.0% BSA, 0.01% Tween-20 for 45 minutes to one hour at room temperature, or overnight at 4°C. Cells were incubated in primary antibody diluted in blocking buffer for two hours, or overnight at 4°C, followed by three 10-minute washes in BSA-PBS. After washing, oocytes were incubated in the secondary antibodies diluted in blocking buffer for one hour, followed by three 10-minute washes in BSA-PBS. DNA was stained using 20µM Hoechst (Thermo Fisher) in BSA-PBS for 15 minutes, and briefly rinsed in BSA-PBS. Oocytes were mounted on no. 1.5 microscope slides (VWR) using ProLong Diamond Antifade Mount (Thermo Fisher) and allowed to dry overnight at room temperature. Slides were stored at 4°C until imaged.

Information on antibodies used in this study can be found in table S3.

Chromosome spreads were allowed to warm to room temperature, then rinsed three times in PBS. Blocking was performed with 3.0% BSA in PBS for 45 minutes. Primary antibodies were diluted in blocking buffer and incubated on the chromosome spreads for two hours, followed by three 15-minute washes with PBS. Secondary antibodies were diluted in blocking buffer and incubated on the spreads for one hour, followed by PBS washes. DNA was then stained using 20µM Hoechst for 15 minutes followed by a PBS rinse. ProLong Diamond was applied to the samples, and a no. 1.5 coverslip was applied and allowed to dry at room temperature. Slides were stored at 4°C until imaged.

### Confocal Microscopy

Immunofluorescence images were obtained at the Optical Imaging and Vital Microscopy Core at Baylor College of Medicine on a Zeiss LSM 780 or 880 Confocal Microscope. Z-stacks were captured using optimal step sizes from bottom of the region of interest to the top of the region. For imaging that was used for fluorescent quantification, parameters were held consistent across all samples.

### Chromosome analysis

Chromosome alignment was analyzed utilizing oocytes cultured for 8.5 hours and fixed as described above. Oocytes were then stained for alpha-tubulin and DNA, then z-stacks of the chromosomes were collected for analysis. Oocytes with the majority of chromosomes aligned at the metaphase plate were classified as ‘good’, while oocytes with multiple chromosomes not localized to the metaphase plate were classified as ‘poor’. To analyze SN and NSN chromosome configuration of GV oocytes, oocytes were collected, denuded, fixed, and DNA was stained with Hoechst. Oocytes with a solid ring of DNA around the nucleolus were classified as SN, while oocytes lacking a ring of DNA around the nucleolus, or containing only a partial ring, were classified as NSN.

### In *vitro* transcription assay

Oocytes were collected and cultured in the presence of 2mM 5-ethynyl uridine (5EU) (Click-IT RNA Alexa Fluor 488 Imaging Kit, Invitrogen) in CZB supplemented with L-glutamine according to the manufacturer’s recommended procedure. In brief, oocytes were cultured in 5EU for 45 minutes then immediately fixed in 3.7% PFA for 20 minutes and rinsed through five drops of BSA-PBS. The oocytes were permeabilized with 0.1% Triton X100 in BSA-PBS for 15 minutes followed by BSA-PBS rinse.

The ClickIT labeling reaction was performed by incubating the oocytes in the provided reaction cocktail according to the manufacturer’s protocol. Oocytes were then counterstained with 20µM Hoechst for 15 minutes, rinsed in BSA-PBS, and mounted in ProLong Diamond, and allowed to dry overnight.

Fluorescent Z-stacks of each oocyte were obtained for analysis. Oocytes containing positive fluorescent signal above the signal of negative control oocytes were considered to be transcriptionally active.

### RNA isolation for qPCR

RNA was isolated from denuded GV oocytes using RNA PicoPure Isolation Kit (Applied Biosystems) and treated with DNase (Qiagen) prior to the generation of cDNA. cDNA was generated using High-Capacity RNA-to-cDNA reverse transcription kit (Invitrogen), and the generated cDNA was subsequently used to analyze gene expression via real-time quantitative PCR (qPCR). qPCR was performed using PowerUpSYBR Green Master Mix (Applied Biosystems) and qPCR MACHINE. Primers used for qPCR are listed in Table S2.

### RNA-sequencing

RNA-seq library was generated according to SMART-seq v4 Ultra low input RNA kit (Takara). In brief, isolated cells were lysed in lysis buffer, 3-SMART-seq CDS primer II and V4 oligonucleotide were added for first stranded cDNA synthesis. cDNA was amplified using PCR Primer II A, and subsequently purified using Ampure XP beads (Beckman). Illumina library was prepared using Nextera XT DNA library preparation kit (Illumina) and sequenced using Illumina Novaseq. Raw counts obtained by mRNA sequencing were filtered, normalized, and differentially analyzed (fold change and FDR-adjusted p-value 0.05) using DESeq2 1.36.0 (*72*). Reads were aligned to the mouse genome version mm39. PCA plot and volcano plots were generated using ggplot2 3.3.5 (*73*) and ggrepel 0.9.1. Heatmap was plotted using pheatmap 1.0.12.

### Statistical analysis

All experiments were performed with a minimum of three independent replicates, with sample sizes indicated in the text or figure legend. Statistical analysis was performed using GraphPad Prism 9. Fertility data, immunofluorescent quantification, and qRT-PCR are displayed by the mean and standard error of the mean (s.e.m.). Student’s t-test was used to determine the difference between two groups, and ANOVA followed by post-hoc testing was used to determine the difference between multiple groups.

ELISA data were log-transformed prior to statistical analysis using Student’s t-test. All t-tests were two tailed. Chromosome alignment, chromosome configuration, and transcriptional activity were analyzed using Fisher’s exact test on raw data. Statistically significance was considered P<0.05.

## Acknowlegements

The authors would like to thank the directors and staff at the following advanced technology core facilities at Baylor College of Medicine: Human Tissue Acqisition and Pathology Core, Optical Imaging and Vital Microscopy Core, and the Single Cell Genomics Core. We additionally thank Dr. Diana Monsivais and Bethany Patton (Baylor College of Medicine) for helpful discussions and comments, and Dr. Cecilia Blengini for assistance with chromosome spreads.

## Author Contributions

Conceptionalization: S.M.B., K.S, and S.A.P. Methodology/Investigation: S.M.B, A.A.A, P.J., and S.A.P. S.M.H. provided the *Ube2i^loxp/loxp^* mice. Funding acquisition: S.A.P., K.S, and S.H. Writing- original draft: S.M.B and S.A.P; Reviewing and editing: S.M.B, S.A.P., K.S., and S.M.H.

## Data and material availability

All sequence data have been deposited in GEO under the accession number, GSE218484. All other data and reagents are available in the Supplemental Materials, and by request to the authors.

