## Supplementary material for "The E2 SUMO-conjugating enzyme UBE2I coordinates the oocyte and zygotic transcriptional programs": Suppl. Figures and Tables

**SUPPLEMENTARY FIGURES**

**Fig. S1**

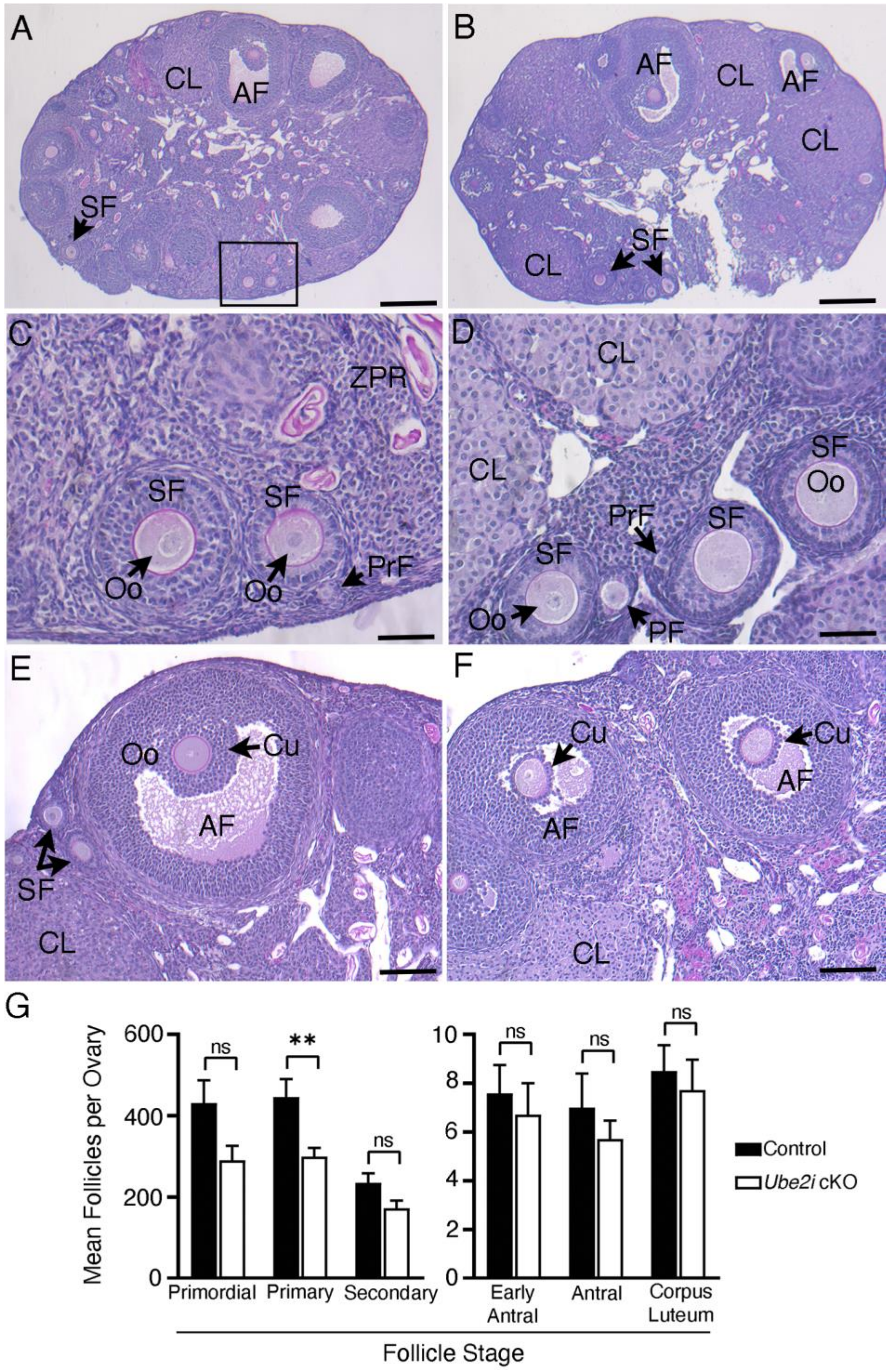


**Fig. S1. Histologic and morphometric analysis of 6-month-old control (A, C, E) and *Ube2i Zp3-cre+* (B, D, F) ovaries shows maintenance of the ovarian reserve with minimal follicle loss**. Ovaries were formalin fixed, processed, and embedded in paraffin, serially sectioned, and stained with periodic acid-Schiff (PAS). (A)

Ovary from a control female showing late stages of folliculogenesis: antral follicles (AF), and corpora lutea (CL). The boxed region contains follicles of earlier stages shown in higher magnification in panel C: secondary follicle (SF), a transitioning primordial/primary follicle (PrF), zona pellucida remnants (ZPR) and oocytes (Oo). (B) *Ube2i Zp3-cre+* ovary contains similar stages. A higher magnification of the conditional knockout ovary is shown in panel D, but from a different animal than in B. (E) Example of a control ovary with a preovulatory antral follicle with cumulus cells (Cu). Cumulus cells in antral follicles of *Ube2i Zp3-cre+* appear to have a less well-developed cumulus cell layer. (G) No difference in follicle stages between genotypes except at the primary follicle stage ($P<0.01$ by t-test), $n=6$ ovaries per genotype. Mean +/- s.e.m. is shown. Scale bars: panels A, B, 400 μm; panel C, D is 50 μm; panels E, F, 100 μm.

**Fig. S2**

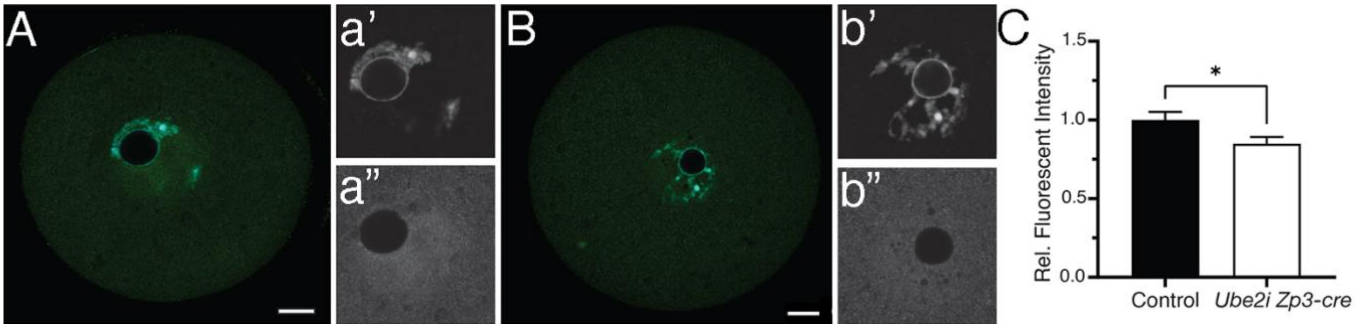


**Fig. S2. Total histone H3.3 protein is significantly decreased in *Ube2i Zp3-cre+* GV oocytes.** GV oocytes were collected and immunostained for histone variant H3.3 (green) and DNA (Hoechst) (blue). (A) Representative z-slice of a control (A) GV oocyte for total histone H3.3 (green), including protein incorporated into the chromosomes and free H3.3. (B) Merged image of z-slices of a *Ube2i Zp3-cre+* oocyte GV for total histone H3.3 (green), including protein incorporated into the chromosomes and free H3.3. Panels to the right of each image shows DNA (a’, b’) and histone H3.3 (a”, b”) staining. C) Relative fluorescence quantification of H3.3 signal. Representative confocal z-slice, scale bars are in A and B are 20µm. (*P = 0.0308, unpaired t-test. Control n = 26, *Ube2i Zp3-cre+* n = 24.)

**Fig S3.**

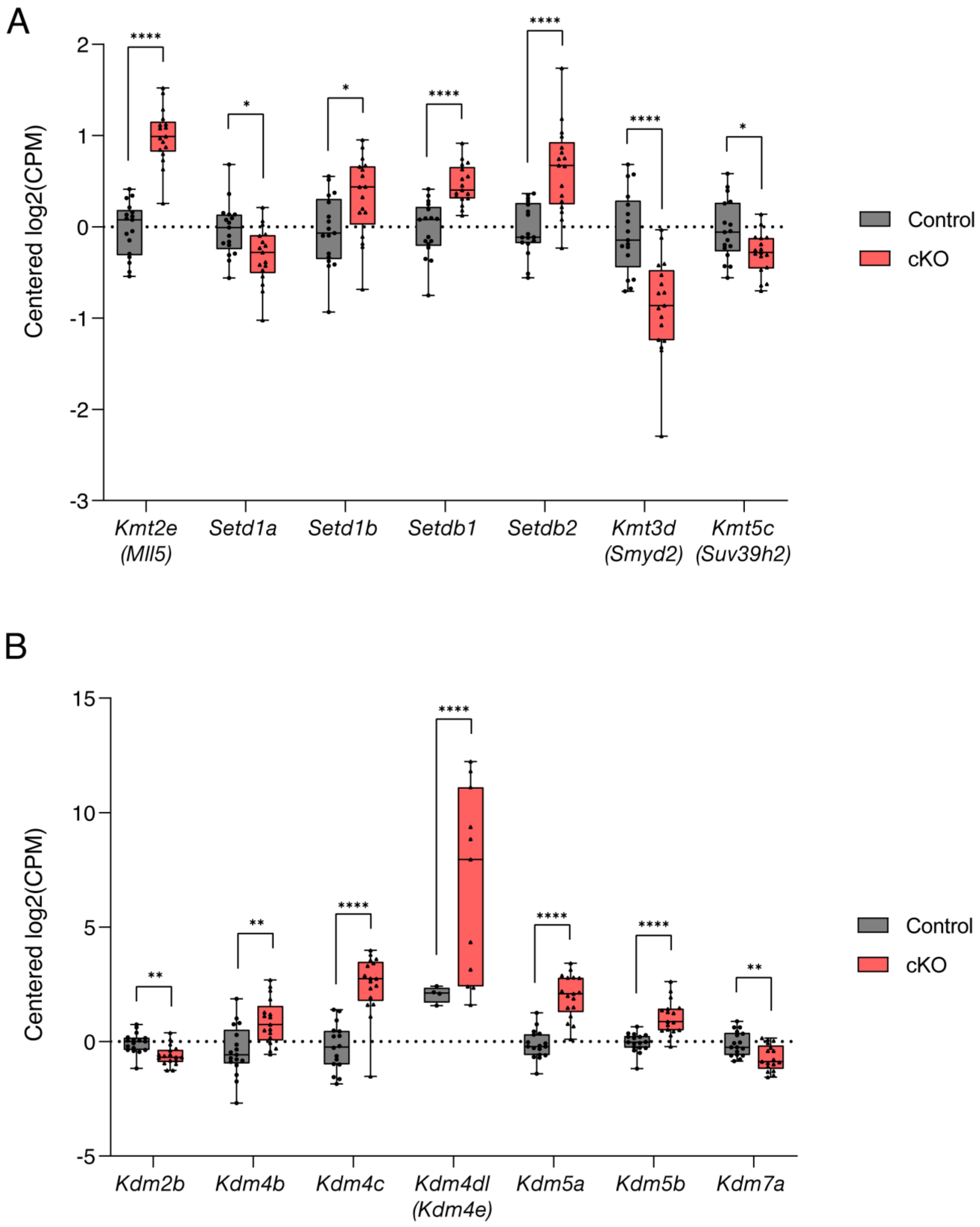
A
Centered log2(CPM)
2
1
0
-1
-2
-3
****
*
*
****
****
****
*
Kmt2e (Mll5)
Setd1a
Setd1b
Setdb1
Setdb2
Kmt3d (Smyd2)
Kmt5c (Suv39h2)
Control
cKO
B
Centered log2(CPM)
15
10
5
0
-5
**
**
****
****
****
****
**
Kdm2b
Kdm4b
Kdm4c
Kdm4dl (Kdm4e)
Kdm5a
Kdm5b
Kdm7a
Control
cKO

**Fig. S3. Differentially expressed histone lysine methyltransferases and demethylases.** A) Histone lysine methyltransferases responsible for trimethylation of H3K4 or H3K9 that are differentially regulated in *Ube2i Zp3-cre+* GV oocytes. B) Demethylases responsible for demethylation of H3K4me3 or H3K9me3 that are differentially regulated in *Ube2i Zp3-cre+* GV oocytes.

**Fig. S4**

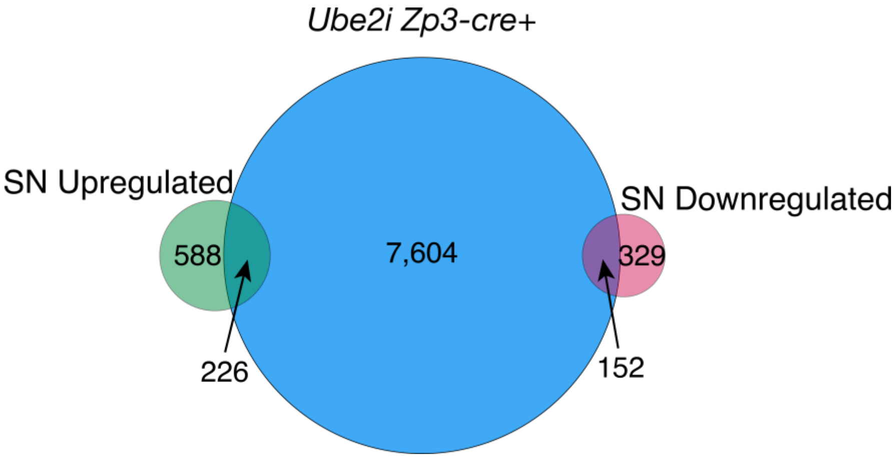


**Fig. S4. DEGs in *Ube2i Zp3-cre+* show no clear correlation with genes associated with the SN or NSN chromosome conformation.** Genes previously reported to be up or downregulated in GV oocytes with a SN chromosome conformation *(28)* were compared to DEGs obtained from RNA-seq analysis of *Ube2i Zp3-cre+* oocytes. No clear correlation between this gene set and *Ube2i Zp3-cre+* DEGs is observed.

**SUPPLEMENTARY TABLES**

**Table S1. Comparison of phenotypes mouse oocyte-specific deletions, *Ube2i Gdf9-icre* and *Ube2i Zp3-cre***

| | ***Ube2i Gdf9-iCre*** | ***Ube2i Zp3-cre*** |
|---|---|---|
| ***Ube2i* deleted region** | Exons 2-3 (null allele) | Exons 3-4 (null allele) |
| ***Cre* expression – cell type** | Oocyte-specific | Oocyte-specific |
| ***Cre* expression - timing** | Primordial follicle to later stages | Primary follicle to later stages |
| **Fertility** | Sterile | Sterile |
| **Superovulation** | Significantly reduced | Significantly reduced |
| **Natural ovulation** | CLs absent | CLs present |
| **Ovarian Reserve** | • Significantly reduced preantral and antral at 8wk<br>• Significantly reduced primordial, preantral, and antral at 14wk<br>• Total depletion at 6mo | Decreased primary follicles at 6mo but maintenance of all other stages |
| **Meiotic Resumption** | Reduced | Reduced |
| **Meiotic Progression** | Arrest at metaphase I | Arrest at metaphase I |
| **Chromosome Alignment** | Not analyzed | Misaligned at metaphase I |
| **Gene Expression** | Altered expression of oocyte-specific genes | Significantly altered transcriptome, premature expression of zygotic genome |

**Table S2. PCR Primers used in this study.**

| Target | Application | Primer | Sequence (5'-3') |
|---|---|---|---|
| *Zp3-cre* | Genotyping | Forward | GCGGTCTGGCAGTAAAAACTATC |
| | | Reverse | GTAAAACAGCATTGCTGTCACTT |
| *Ube2i* 5' *loxP* site | Genotyping | Forward | AGGTAGGGGTGGCTTAGAGG |
| | | Reverse | GGTTCATTGTGCCATCAGGG |
| *Ube2i 3'* *loxP* site | Genotyping | Forward | CAAGTCCCAGGGTAGATGCG |
| | | Reverse | CAGCTCAGACCTGGCCTTAC |
| *Duxf3* | qPCR | Forward | AACCCACGACCAGGCTTTG |
| | | Reverse | CCGAGCTCTTCGGTTTTGAA |
| *Eif1a* | qPCR | Forward | AAAAACAGGCGCAGAGGTAAA |
| | | Reverse | TCCTCACACCGTCAAAGCAC |
| *MuERV-L* | qPCR | Forward | ATCTCCTGGCACCTGGTATG |
| | | Reverse | AGAAGAAGGCATTTGCCAGA |
| *Tmem92* | qPCR | Forward | GGGGCACACTCACCTTGAC |
| | | Reverse | CAGCATTCCTTGACACAGCAT |
| *Ube2i* | qPCR | Forward | AAAGGAAAGCCTGGAGGAAG |
| | | Reverse | CTCCCAGTTCATCAGGTTCAT |
| *Zscan4* | qPCR | Forward | GAGATTCATGGAGAGTCTGACTGATGAGTG |
| | | Reverse | GCTGTTGTTTCAAAAGCTTGATGACTTC |

**Table S3. Antibodies used in this study.**

| Antibody | Species | Vender | Catalog No. | Dilution |
|---|---|---|---|---|
| Alexa Fluor 488-α tubulin | Mouse | Invitrogen | 322588 | 1:200 |
| α-Centromere Protein | Human | Antibodies Incorporated | 15-2105 | 1:100 |
| α-H3K4me3 | Rabbit | Cell Signaling Technology | 9751T | 1:300 |
| α-H3K9me3 | Rabbit | Cell Signaling Technology | 13969T | 1:300 |
| α-H3.3 | Rabbit | Abcam | Ab176840 | 1:100 |
| α-Human Alexa Fluor 647 | Goat | Invitrogen | A21445 | 1:250 |
| α-Rabbit Alexa Fluor Plus 488 | Goat | Invitrogen | A32731 | 1:250 |
| α-Rabbit Alexa Fluor Plus 594 | Goat | Invitrogen | A32740 | 1:250 |
